# A dual endosymbiosis drives nutritional adaptation to hematophagy in the invasive tick *Hyalomma marginatum*

**DOI:** 10.1101/2021.10.12.464044

**Authors:** Marie Buysse, Anna Maria Floriano, Yuval Gottlieb, Tiago Nardi, Francesco Comandatore, Emanuela Olivieri, Alessia Giannetto, Ana Palomar, Ben Makepeace, Chiara Bazzocchi, Alessandra Cafiso, Davide Sassera, Olivier Duron

**Author notes:** Co-first authors (Marie Buysse, Anna Maria Floriano). Co-lead authors (Davide Sassera, Olivier Duron).

## Abstract

Many animals are dependent on microbial partners that provide essential nutrients lacking from their diet. Ticks, whose diet consists exclusively on vertebrate blood, rely on maternally inherited bacterial symbionts to supply B vitamins. While previously studied tick species consistently harbor a single lineage of those nutritional symbionts, we evidence here that the invasive tick *Hyalomma marginatum* harbors a unique dual-partner nutritional system between an ancestral symbiont, *Francisella*, and a more recently acquired symbiont, *Midichloria*. Using metagenomics, we show that *Francisella* exhibits extensive genome erosion that endangers the nutritional symbiotic interactions: Its genome includes folate and riboflavin biosynthesis pathways but deprived functional biotin biosynthesis on account of massive pseudogenization. Co-symbiosis compensates this deficiency since the *Midichloria* genome encompasses an intact biotin operon, which was primarily acquired via lateral gene transfer from unrelated intracellular bacteria commonly infecting arthropods. Thus, in *H. marginatum*, a mosaic of co-evolved symbionts incorporating gene combinations of distant phylogenetic origins emerged to prevent the collapse of an ancestral nutritional symbiosis. Such dual endosymbiosis was never reported in other blood feeders but was recently documented in agricultural pests feeding on plant sap, suggesting that it may be a key mechanism for advanced adaptation of arthropods to specialized diets.

## Introduction

Ticks evolved to hematophagy through the acquisition of key genomic adaptations (Gulia-Nuss et al., 2016; Jia et al., 2020) and the evolution of mutualistic interactions with microbial symbionts (Buysse and Duron, 2021; Duron and Gottlieb, 2020). Ticks rely on vertebrate blood as their sole food source throughout their development, although it is nutritionally unbalanced: Blood contains high levels of proteins, iron, and salts, but low levels of carbohydrates, lipids, and vitamins. To overcome these dietary challenges, tick genomes present large repertoires of genes related to vitamin and lipid shortages, iron management, and osmotic homeostasis (Gulia-Nuss et al., 2016; Jia et al., 2020). Yet, tick genomes do not contain genes for synthesis of some essential vitamins, thus vitamin-provisioning pathways have indirectly arisen from symbiotic associations with transovarially transmitted intracellular bacteria (Ben-Yosef et al., 2020; Duron et al., 2018; Duron and Gottlieb, 2020; Gerhart et al., 2016; Gottlieb et al., 2015; Guizzo et al., 2017; Smith et al., 2015). This partnership allowed the long-term acquisition of heritable biochemical functions by ticks and favored their radiation into the nearly 900 tick species that currently exist (Duron and Gottlieb, 2020). This co-evolutionary process is reflected by microbial communities dominated by tick nutritional symbionts (Bonnet and Pollet, 2020; Narasimhan and Fikrig, 2015).

Most tick species typically harbor a single lineage of nutritional symbiont, but multiple symbiont lineages exist in different tick species (Duron and Gottlieb, 2020). The most common lineages belong to the *Coxiella*-like endosymbiont group and to the *Francisella*-like endosymbiont group (CLE and FLE, hereafter) (Azagi et al., 2017; Binetruy et al., 2020; Duron et al., 2017; Machado-Ferreira et al., 2016), but a few tick species may rely on other symbionts (Duron et al., 2017), such as the intramitochondrial bacterium *Midichloria* (Sassera et al., 2006). Once deprived of their nutritional symbionts, ticks show a complete stop of growth and moulting, as well as lethal physical abnormalities (Ben-Yosef et al., 2020; Duron et al., 2018; Guizzo et al., 2017; Zhong et al., 2007), which can be fully restored with an artificial supplement of B vitamins (Duron et al., 2018). Genomes of CLE and FLE consistently contain complete biosynthesis pathways of the same three B vitamins: biotin (vitamin B_7_), riboflavin (B_2_), and folate (B_9_) (Duron et al., 2018; Gerhart et al., 2018, 2016; Gottlieb et al., 2015; Guizzo et al., 2017; Nardi et al., 2021; Smith et al., 2015). These three pathways form a set of core symbiotic genes essential for ticks (Duron and Gottlieb, 2020) while other functions are disposable: As such, the genomes of CLE, FLE and *Midichloria* underwent massive reduction as observed in other nutritional endosymbionts of arthropods with a specialized diet (Duron et al., 2018; Gerhart et al., 2018, 2016; Gottlieb et al., 2015; Nardi et al., 2021; Sassera et al., 2011; Smith et al., 2015). This reduction process is primarily initiated as a consequence of relaxed selection on multiple genes which are not essential within the host stable and nutrient-rich cellular environment (Bennett and Moran, 2015; McCutcheon et al., 2019). Over time, genes encoding for functions such as motility, regulation, secondary-metabolite biosynthesis, and defense are pseudogenized and then definitively jettisoned (Bennett and Moran, 2015; McCutcheon et al., 2019).

Coevolution of hosts and their nutritional symbionts over millions of years is expected to result in stable co-cladogenesis (Bennett and Moran, 2015). However, in sap-feeding insects where symbionts compensate for nutritional deficiencies of the sap diet, cases of unstable interactions have been observed (Bennett and Moran, 2015; McCutcheon et al., 2019). Indeed, as the symbiont genome decays, repair mechanisms can be lost, leading to faster pseudogenization, which can affect genes in all functional categories including those involved in nutritional supplementation (Bennett and Moran, 2015; McCutcheon et al., 2019). This may ultimately lead to overly reduced and maladapted genomes, limiting beneficial contributions to hosts and opening the road to wholesale replacement by new symbiont(s) (Bennett and Moran, 2015; McCutcheon et al., 2019; Sudakaran et al., 2017). This degeneration–replacement model has been proposed in sap-feeding insects (Campbell et al., 2015; Łukasik et al., 2017; Manzano-Marín et al., 2020; Matsuura et al., 2018), but is difficult to document as such replacements are expected to be transient (McCutcheon et al., 2019). In parallel, other forms of multi-partite interactions have been reported, where a novel bacterium with similar function to a pre-existing symbiont enters a bipartite system (Husnik and Mccutcheon, 2016; Santos-Garcia et al., 2018; Takeshita et al., 2019). This can lead to relaxation of conservative selection pressure on the original symbionts, which can lose genes that were previously fundamental, ultimately leading to a tripartite stable symbiosis (Husnik and Mccutcheon, 2016; Takeshita et al., 2019).

In this study we document a complex nutritional endosymbiosis in the invasive bont-legged tick *Hyalomma marginatum* characterized by a maladapted ancestral symbiont genome compensated by gene functions of a more recently acquired symbiont. This large tick species usually feeds on humans and a diversity of mammals and birds in the Mediterranean climatic zone but has recently expanded its range to the north, invading Western Europe (Vial et al., 2016). *Hyalomma marginatum* can transmits Crimean-Congo hemorrhagic fever virus, an emerging disease affecting humans in the northern Mediterranean Basin, as well as protozoan parasites and intracellular bacterial pathogens such as *Rickettsia aeschlimannii* (Vial et al., 2016). Previous studies detected the presence of diverse highly prevalent bacterial symbionts across *H. marginatum* populations (Di Lecce et al., 2018), but their contribution to tick nutrition was not studied. Here we sequenced three microbial metagenomes of *H. marginatum* in three distinct locations, and determined the nutritional interactions in this unique symbiosis system.

## Results

### Microbial metagenomes

We sequenced three microbial metagenomes from *H. marginatum* specimens collected in Spain (i.e., the Hmar-ES metagenome), Italy (Hmar-IT), and Israel (Hmar-IL). We obtained 57,592,510 paired-end reads from the Hmar-ES metagenome, 68,515,944 paired-end reads from Hmar-IT, and 10,807,740 paired-end reads from Hmar-IL. We assembled one complete FLE genome from each metagenome (FLE-Hmar-ES, FLE-Hmar-IT, and FLE-Hmar-IL; also designated as FLE-Hmar genomes). In parallel, we assembled one complete genome from each metagenome for the bacterium *Midichloria*, and thus obtained three additional complete genomes (Midi-Hmar-ES, Midi-Hmar-IT, and Midi-Hmar-IL; also designated as Midi-Hmar genomes). In addition, we detected a third bacterium, the tick-borne pathogen *Rickettsia aeschlimannii*, in the Hmar-IT and Hmar-IL, but not Hmar-ES, metagenomes, and further assembled one complete (RAES-Hmar-IT) and one partial (RAES-Hmar-IL) genome (Figure 1).

**Figure 1.**
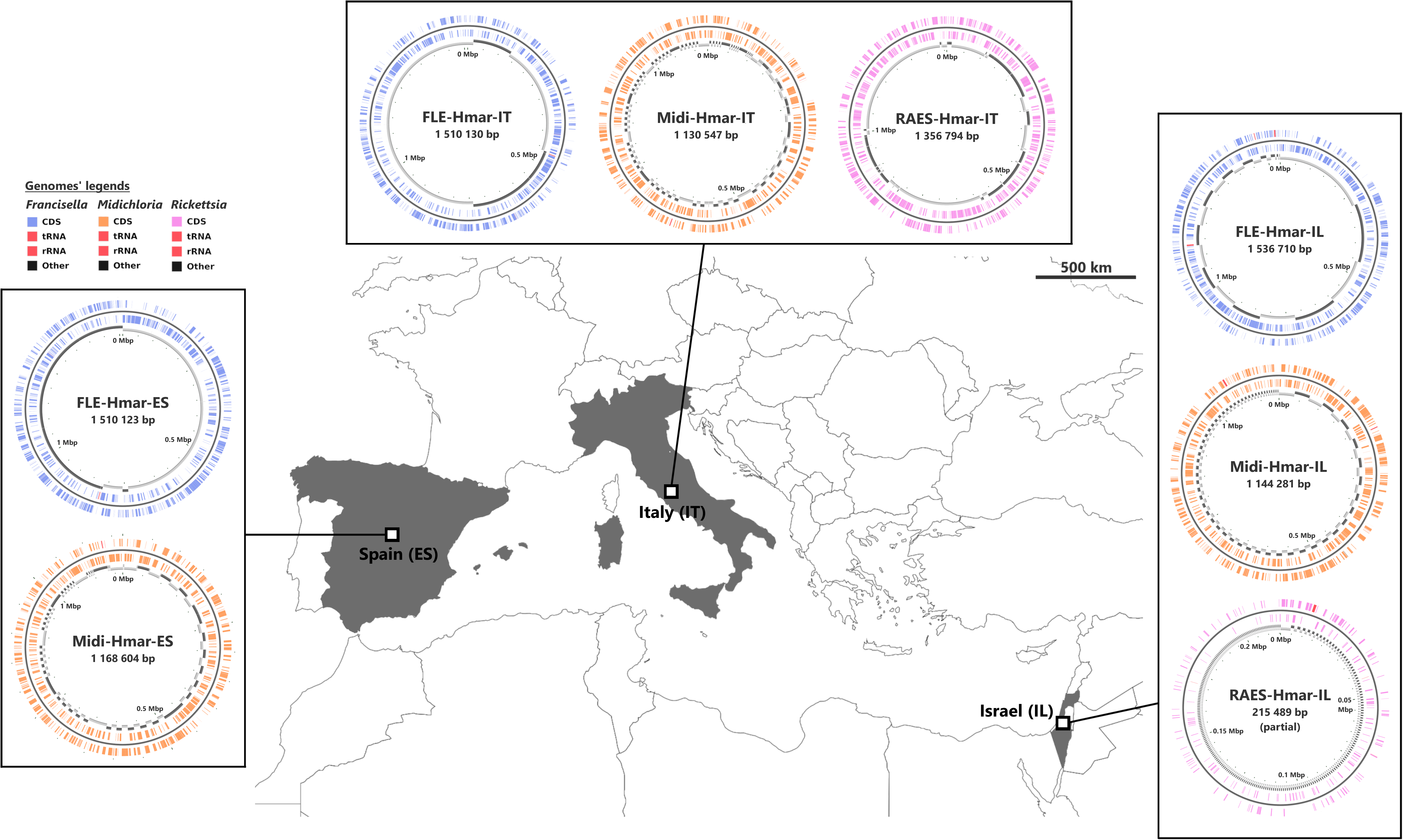
Sampled locations for *H. marginatum* ticks. Sampling countries are colored in grey. The symbiont genomes retrieved from each site are graphically represented in blue, orange and pink, corresponding to *Francisella*-LE (FLE), *Midichloria* (Midi) and *R. aeschlimannii* (RAES), respectively. Circles on genome maps correspond to the following: (1) forward strand genes; (2) reverse strand genes, (3) in grey and black, contigs. Circular maps of the genomes were produced using CGView.

The assemblies of the three FLE draft genomes (13 contigs for FLE Hmar-IT, 17 for FLE Hmar-ES, and 25 FLE Hmar-IL) were very similar in size (1.51 to 1.536 Mb), average G+C content (31.11 to 31.23%), number of predicted protein-coding genes (937 to 961), and protein-coding sequences with inactivating mutations (pseudogenes, 789 to 798), but no signal of mobile elements (Supplementary Table S1). Phylogenomic analysis based on 436 single-copy orthologous *Francisella* genes showed that FLE-Hmar-ES, FLE-Hmar-IT, and FLE-Hmar-IL cluster together in a robust clade within other FLE of ticks (Figure 2A). The closest relative of this clade is the FLE of another *Hyalomma* species, *Hyalomma asiaticum* from China, while FLEs of other tick genera (*Ornithodoros*, *Argas*, and *Amblyomma*) are more distantly related. The FLE clade is embedded in the *Francisella* genus and shares a common origin with pathogenic species, including the tularemia agent, *F. tularensis*, and an opportunistic human pathogen, *F. opportunistica* (Figure 2A). The FLE-Hmar genomes are similar in organization and content overall, apart from a 127,168 bp inversion in FLE-Hmar-ES and a 4,165 bp inversion in FLE-Hmar-IL (Figure 2B). Compared to other FLE genomes (i.e. FLEOm and FLEAm), they are slightly reduced in size and exhibit substantial rearrangements (Figure 2B, Tables S1 and S2). Based on gene orthologs, the core-genome of these FLE shares 743 genes, while the core genome of the FLE of *H. marginatum* contains 796 genes because the FLE-Hmar genomes share 53 genes that are not present in other FLE genomes (Figure 2C). Interestingly, pseudogenes are highly abundant in the three genomes of FLE of *H. marginatum,* compared with other FLE genomes (Figure 2D), and most functional categories are impacted by this advanced pseudogenization process. Most of the genes associated with virulence in pathogenic *Francisella* species, including the *Francisella* Pathogenicity Island (FPI) and secretion system, are pseudogenized or completely missing in FLE-Hmar genomes, consistent with previous observations in other FLEs (Figure S1). Furthermore, FLE-Hmar genomes contain fewer genes involved in replication, recombination, and repair than any other FLEs analyzed (Figure 2C). The nucleotide excision repair system (*uvrABCD*) in FLE-Hmar genomes is conserved, however the DNA mismatch repair system (*mutSL*) is fully pseudogenized. In other FLEs, these systems are conserved, with the exception of the *mutSL* system in the FLE of *Ornithodoros moubata* and *H. asiaticum* genomes (Figure S2).

**Figure 2.**
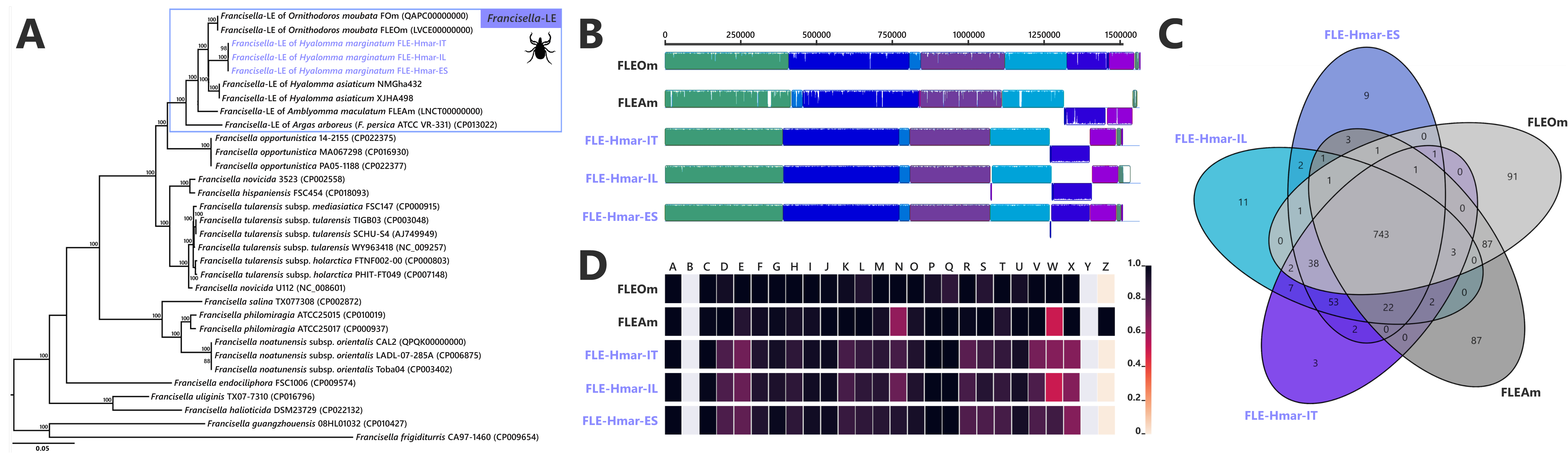
Comparative genomic features of FLE. (A) Whole genome phylogenetic relationship of the three FLE genomes obtained from *H. marginatum* specimens (highlighted), FLE of other tick species, and *Francisella* pathogenic species, including *F. tularensis*. The phylogenetic tree was inferred using maximum likelihood from a concatenated alignment of 436 single-copy orthologous genes (132,718 amino acids; best-fit approximation for the evolutionary model: CPREV+G4+I). The numbers on each node represent the support of 1,000 bootstrap replicates; only bootstrap values >70% are shown. The scale bar is in units of substitution/site. Sequences from newly sequenced FLE genomes are indicated by a blue font. (B) Whole genome synteny of the three FLE genomes obtained from *H. marginatum* specimens (blue font) and representative FLE (black font). Each contiguously colored locally collinear block (LCB) represents a region without rearrangement of the homologous backbone sequence. LCBs below the center in *Francisella* genomes represent blocks in the reverse orientation. The height of color bars within LCBs corresponds to the average level of similarity (conservation) profile in that region of the genome sequence. Areas that are completely white were not aligned and contain sequence elements specific to a particular genome. (C) Venn diagram representing orthologs distribution between the three FLE genomes obtained from *H. marginatum* specimens (blue font) and representative FLE (black font). (D) Clusters of orthologous groups of proteins (COGs) comparisons between the three FLE genomes obtained from *H. marginatum* specimens (blue font) and representative FLE (black font). The color scale indicates the percentage of functional genes per category. Grey color means that genes are not present in a given category. The COG category names (capital letters from A to Z) correspond to the NCBI reference naming system.

The *Midichloria* genomes, Midi-Hmar-ES, Midi-Hmar-IT, and Midi-Hmar-IL, were assembled in 179, 359 and 123 contigs respectively, and their genomes reach 1.13 to 1.16 Mb, with consistent G+C content (35.03-35.23%), number of predicted protein-coding genes (1,032 to 1,071), pseudogenes (190 to 192), and no signal of mobile elements (Table S1). Remarkably, their average coding density (76%) is higher than those of FLEs of *H. marginatum* (54%), and furthermore, those of other FLE representatives (range from 57 to 69%). Phylogenomic analysis based on 242 single-copy orthologous *Rickettsiales* genes shows that Midi-Hmar-ES, Midi-Hmar-IT, and Midi-Hmar-IL cluster in a highly supported monophyletic sister group of the single other sequenced *Midichloria* genome, *Candidatus* M. mitochondrii strain IricVA (*M. mitochondrii* hereafter), the intra-mitochondrial endosymbiont of the European sheep tick, *Ixodes ricinus*. Altogether, they delineate a monophyletic clade within the *Midichloriaceae* family (Figure 3A). The high number of Midi-Hmar contigs precluded comparison of their genome structures. However, gene contents of the Midi-Hmar genomes are highly similar, differing substantially from *M. mitochondrii* (Tables S1 and S2). Based on gene orthologs, the pan-genome of *Midichloria* shared 745 genes, and Midi-Hmar genomes shared 181 genes that are not present in *M. mitochondrii* (Figure 3B). Pseudogenization only impacted a few functional categories (e.g., genes of chromatin structure and dynamics [category B] and genes of the mobilome [prophages and transposons, category X]), across Midi-Hmar genomes (Figure 3C). By contrast to FLE genomes, the DNA mismatch repair system (*mutSL*) and the nucleotide excision repair (*uvrABCD*) are not pseudogenized in Midi-Hmar genomes and are thus potentially functional (Figure S2).

**Figure 3.**
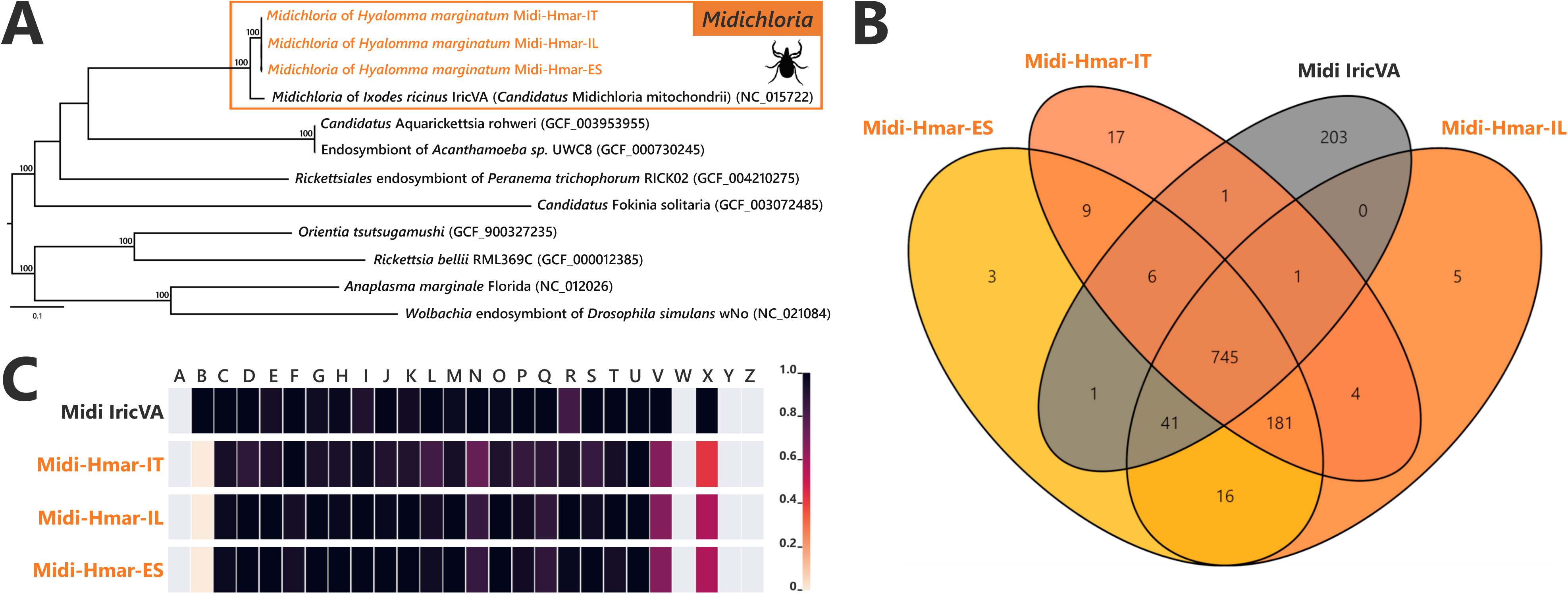
Comparative genomic features of *Midichloria*. (A) Whole genome phylogenetic relationship of the three *Midichloria* genomes obtained from *H. marginatum* specimens (highlighted), *M. mitochondrii*, and other *Rickettsiales* species. The phylogenetic tree was inferred using maximum likelihood from a concatenated alignment of 242 single-copy orthologous genes (49,957 amino acids; best-fit approximation for the evolutionary model: LG+G4+I). The numbers on each node represent the support of 1,000 bootstrap replicates; only bootstrap values >70% are shown. The scale bar is in units of substitution/site. Sequences from newly sequenced *Midichloria* genomes are indicated by an orange font. (B) Venn diagram representing orthologs distribution between the three *Midichloria* genomes obtained from *H. marginatum* specimens (orange font) and “*Ca*. M. mitochondrii” (black font). (C) Clusters of Orthologous Groups of proteins (COGs) comparisons between the three *Midichloria* genomes obtained from *H. marginatum* specimens (orange font) and “*Ca*. M. mitochondrii” (black font). The color scale indicates the percentage of functional genes per category. Grey color means that genes are not present in a given category. The COG category names (capital letters from A to Z) correspond to the NCBI reference naming system.

In two metagenomes, we also detected contigs belonging to *R. aeschlimannii*. We recovered one complete chromosome (1.35 Mb for 165 contigs, with 1,122 genes, 475 pseudogenes and 32.32% G+C content) for RAES-Hmar-IT (Table S1), whereas for RAES-Hmar-IL only a partial genome (0.215 Mb) was obtained. No *Rickettsia* reads were detected in the Hmar-ES metagenome. This may be due to the absence of the bacterium in the Hmar-ES microbiome, or, possibly, to experimental artefact: This metagenome was obtained by sequencing DNA extracted from the dissected ovaries of 10 females, whereas the Hmar-IT metagenome was sequenced from DNA extracted from a whole female, suggesting that *Rickettsia* may be absent or at low density in tick ovaries. This pattern of tissue localization may be corroborated by the Hmar-IL metagenome, sequenced from a pool of dissected ovaries of 10 ticks, but for which we obtained only a few *Rickettsia* reads. This reflects either the absence of *Rickettsia* in the dissected ticks, or a lower abundance of *Rickettsia* in ovaries. Phylogenetic analysis based on 324 single-copy orthologous genes present in *Rickettsia* genomes showed that RAES-Hmar-IT clusters with the tick-associated human pathogen *R. aeschlimannii* (including the strain MC16 previously isolated from *H. marginatum*) and altogether delineate a robust clade with the Spotted Fever *Rickettsia* group (Figure S3). Overall, the low abundance of *Rickettsia* in the Hmar-IL metagenome (8x average genome coverage), its absence in the Hmar-ES metagenome, as well as the genetic proximity with *R. aeschlimannii,* are all indicative of a tick-borne pathogen, rather than a nutritional endosymbiont. The RAES-Hmar-IT and RAES-Hmar-IL assemblies were thus not used for further analyses.

### Complementation in nutritional endosymbiosis

While all previously sequenced FLE genomes contain complete pathways for biotin, riboflavin, and folate, the FLE-Hmar genomes only retain intact pathways for riboflavin and folate (Figure 4A). The biosynthesis pathway for biotin is non-functional, as from six genes of the complete pathway, one is missing and four are pseudogenized (frameshifted) in the FLE-Hmar genomes (Figure 4A). Partial biosynthesis pathways of four other B vitamins (pantothenic acid: B_5_; nicotinic acid: B_3_; pyridoxine: B_6;_ and thiamine: B_1_), each containing one to four pseudo/missing genes, are also present in the FLE-Hmar genomes; a pattern consistent with other FLE genomes (Figure S4).

**Figure 4.**
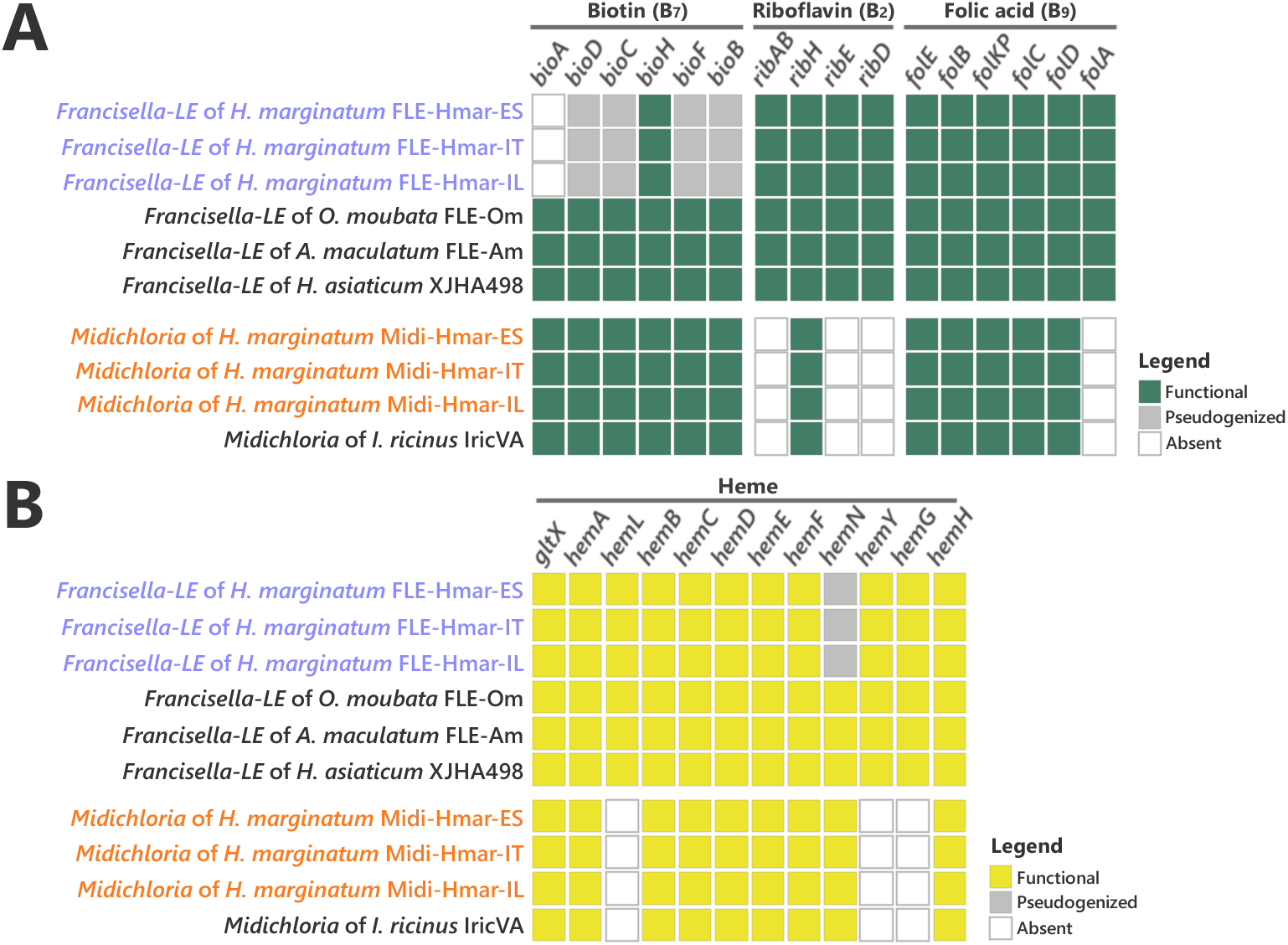
Conservation level of biosynthetic pathways involved in the symbiotic interaction between ticks and endosymbionts. (A) Biosynthetic pathways for the three key B vitamins (biotin, folic acid, and riboflavin) required for tick nutrition of FLE, *F. tularensis*, and *Midichloria*. Newly sequenced FLE and *Midichloria* genomes are indicated by a blue and an orange font, respectively. Green squares, functional genes; grey squares, pseudogenes; white squares, missing genes. (B) Heme biosynthetic pathway of FLE, *F. tularensis*, and *Midichloria*. Newly sequenced FLE and *Midichloria* genomes are indicated by a blue and an orange font, respectively. Yellow squares, functional genes; grey squares, pseudogenes; white squares, missing genes.

The B-vitamin pathways in Midi-Hmar genomes are similar to those of *M. mitochondrii*. A complete pathway of biotin was consistently recovered in *Midichloria* genomes (Figure 4A). However, only partial pathways were detected for folate, riboflavin, and pantothenic acid with one, three, and four missing genes, respectively (Figures 4A and S4).

Ticks differ from other eukaryotes in that they lack most of the genes encoding proteins required to produce and degrade heme (Gulia-Nuss et al., 2016; Jia et al., 2020). We thus searched for relevant pathways in the symbiont genomes and found that the FLE-Hmar and Midi-Hmar genomes contain genes involved in heme and iron homeostasis. The heme biosynthetic pathway was consistently detected, as well as a gene encoding heme oxygenase that catalyzes the degradation of heme and the subsequent release of iron (Figure 4B). In these genomes we detected similar genetic patterns as those observed in the other FLE and *Midichloria* genomes. However, the heme pathway is partially degraded in FLE-Hmar genomes since their *hemN* gene is pseudogenized, a pattern not observed in other FLE genomes, suggesting an early stage of degradation for the heme pathway. This gene is present and not pseudogenized in the Midi-Hmar genomes (Figure 4B), suggesting these symbionts may compensate for the FLE heme pathway. In addition, three other genes, *hemL*, *hemY,* and *hemG* were not found in *Midichloria* genomes, but were present in FLE genomes (Figure 4B). As a result, a combination of FLE and *Midichloria*genes may be required to obtain a complete heme pathway.

### Phylogenetic origins of B vitamin biosynthetic pathways

Our analysis shows that the B vitamin biosynthesis genes in the FLE-Hmar genomes have all been vertically inherited from a *Francisella* ancestor as their B vitamin genes are always more closely related to *Francisella* orthologous genes than to genes of more distantly related bacteria (examples are shown in Figures S5A-B). Similarly, folate and riboflavin biosynthesis genes in the Midi-Hmar genomes are of *Midichloria* origin as these genes are more closely related to orthologous genes present in *M. mitochondrii* and other members of the *Midichloriaceae* family than to genes of other bacteria (Figures S5A-B).

By contrast, the six biotin genes present in the Midi-Hmar genomes did not originate from the *Midichloriaceae* family, but from an exogenous origin. The biotin synthesis genes *bioD*, *bioC*, *bioH*, *bioF,* and *bioB* formed a compact operon in the Midi-Hmar genomes while the *bioA* gene is more distantly located along the bacterial chromosome (Figure 5). A similar operon structure is also present on the *M. mitochondrii* genome. In contrast, an arrangement in which the *bioA* is gene physically associated with the remainder of the biosynthetic operon was identified on the genomes of multiple intracellular bacteria that are all distantly related to *Midichloria,* including the *Cardinium* endosymbionts (Bacteroidetes: Flexibacteraceae) of parasitoid wasps and leaf-hoppers; the *Wolbachia* endosymbionts (Alphaproteobacteria: Anaplasmataceae) of bed bugs, bees, planthoppers, and fleas; the *Legionella* endosymbiont (Gammaproteobacteria: Legionellaceae) of the rodent lice; *Rickettsia buchneri* (Alphaproteobacteria: Rickettsiaceae) symbiont of the blacklegged tick *Ixodes scapularis*; and diverse intracellular pathogens of vertebrates, such as *Lawsonia* (Deltaproteobacteria: Desulfovibrionaceae) and *Neorickettsia* (Alphaproteobacteria: Anaplasmataceae) (Figure 5 and Table S3). In most cases, the biotin genes form a compact and streamlined operon of 5-7 kb without major structural variation among bacteria. Phylogenetic analyses of the six biotin synthesis genes consistently exhibited similar evolutionary patterns (Figure 5). These patterns suggest that the biotin synthesis genes were acquired as a whole operon by an ancestor of *Midichloria* from an unrelated intracellular bacterium. In some cases, this streamlined biotin operon is also associated with nutritional symbionts of obligate blood feeders, such as *Wolbachia* of bedbugs, *Legionella* of rodent lice, and *R. buchneri* in the blacklegged tick, *I. scapularis* (Figure 5).

**Figure 5.**
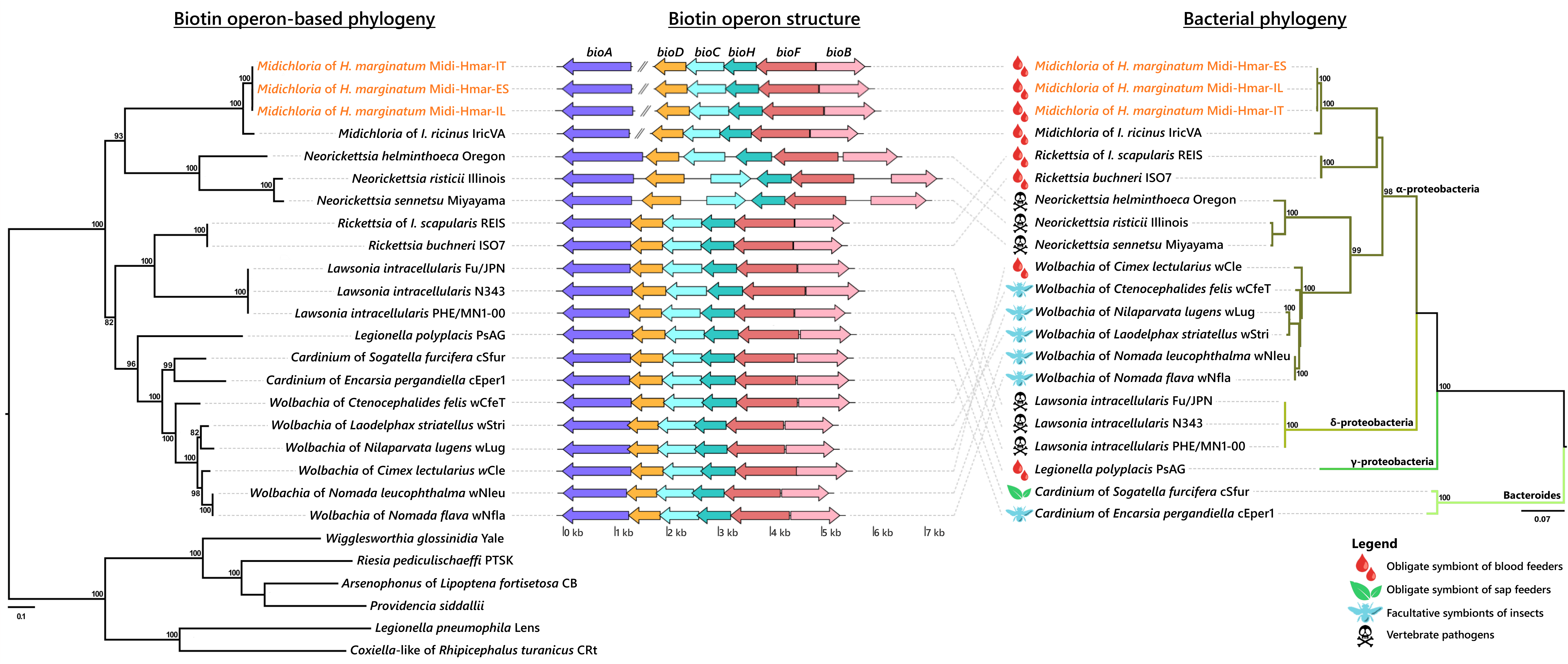
Evolutionary relationships (left) and structure (middle) of the streamlined biotin operon, confronted to the bacterial phylogeny (right). The phylogenetic tree of the streamlined biotin operon (left) was inferred using maximum likelihood from a concatenated alignment of the six genes composing the biotin biosynthetic pathway (1,493 amino acids; best-fit approximation for the evolutionary model: CPREV+G4+I). The numbers on each node represent the support of 1,000 bootstrap replicates; only bootstrap values >70% are shown. The scale bar is in units of substitution/site. Sequences from newly sequenced *Midichloria* genomes are indicated by an orange font. Structure of the streamlined biotin operon (middle) is represented by arrows in which each colored arrow corresponds to an intact gene and its direction. A discontinuous line corresponds to genes encoding on two separated contigs. The bacterial phylogenetic tree (right) was inferred using maximum likelihood from an alignment of 16S rDNA sequences (1,362 bp unambiguously aligned; best-fit approximation for the evolutionary model: GTR+G+I). The numbers on each node represent the support of 1,000 bootstrap replicates; only bootstrap values >70% are shown. The scale bar is in units of substitution/site. Sequences from *Midichloria* of *H. marginatum* are indicated by an orange font.

### Co-infections of nutritional endosymbionts in tick ovaries

Real-time qPCR assays on 10 field *H. marginatum* adult females indicated that FLE and *Midichloria* are both abundant in tick ovaries, with FLE and *Midichloria* detected in 10 and 9 ovarian samples, respectively. High densities were always observed for FLE (686.09 ± 206.89 FLE *rpoB* gene copies per tick *cal* gene copies) and for *Midichloria* (88.72 ± 28.41 *Midichloria gyrB* gene copies per tick *cal* gene copies) (Figure 6A). FLE were consistently most abundant with densities 5.8 to 21.4 × higher than *Midichloria* densities (paired t test, *p* = 0.013; one outlier was removed from analysis, n = 8). The FLE and *Midichloria* densities covaried positively in individual ticks (χ2 = 23.937, *p* = 0.003; Figure 6B).

**Figure 6.**
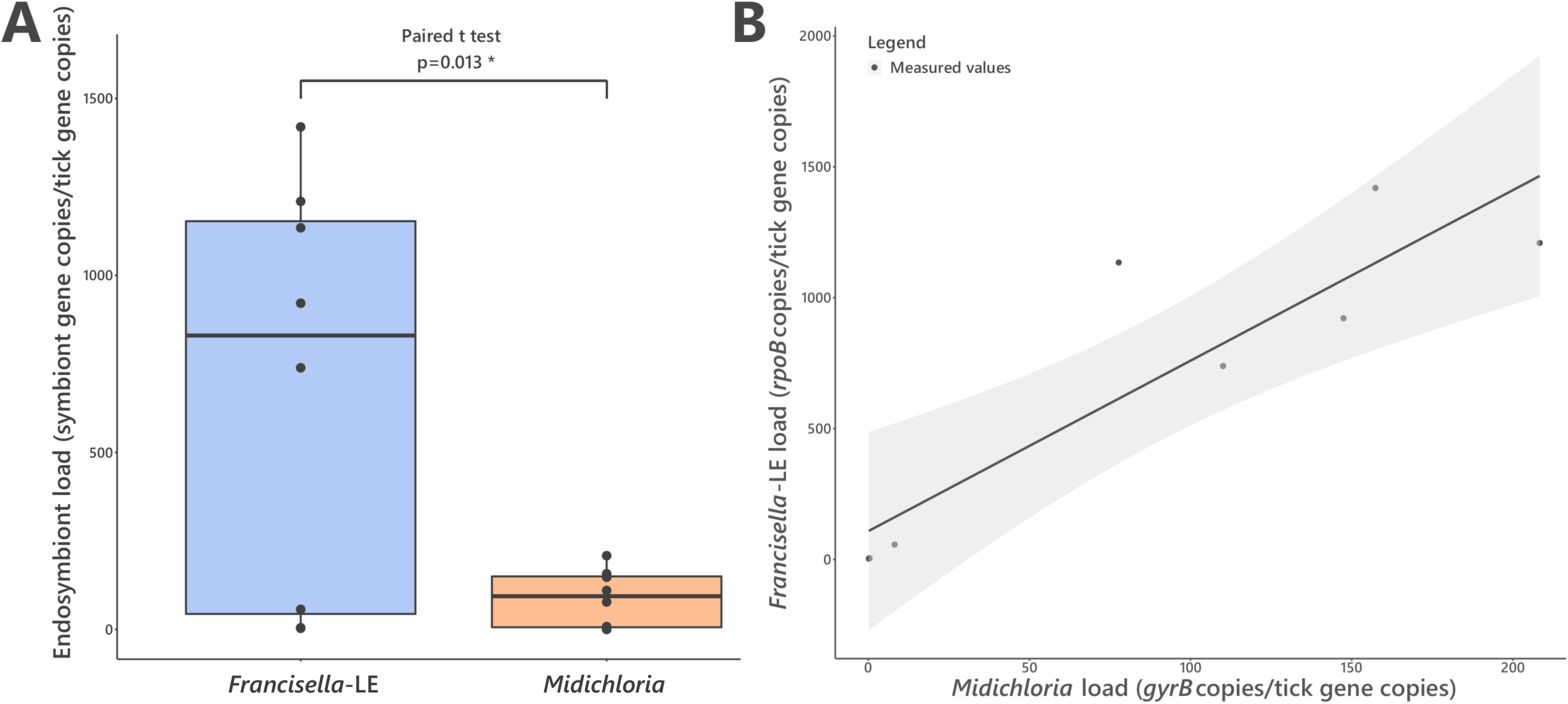
FLE and *Midichloria* loads in ovaries of *H. marginatum*. (A) Representation and statistical comparison of FLE and *Midichloria* densities. (B) FLE density as a function of *Midichloria* density in *H. marginatum* ovaries. Grey area represents 95% confidence interval.

### Evolutionary history of endosymbiosis in the *Hyalomma* genus

We further examined the evolutionary history of *Hyalomma* species and their associations with FLE and *Midichloria* through a bibliographic survey. Of 10 *Hyalomma* species with sufficient published data, all were found infected by FLE, and 9 by *Midichloria* (Table S4). We next reconstructed *Hyalomma* phylogeny using mitochondrial DNA sequences of the cytochrome c oxidase I (*cox1*) gene available on Genbank (Figure S6). Mapping of FLE and *Midichloria* onto the *Hyalomma* phylogenetic tree showed that co-infection by FLE and *Midichloria* is widespread across all *Hyalomma* lineages (Figure S7). However, FLE and *Midichloria* co-infection was not detected in one species, *H. asiaticum*, nested within the *Hyalomma* phylogeny among co-infected species.

## Discussion

In this study, we show that the tick *H. marginatum* is a complex symbiotic system, combining genes of diverse phylogenetic origins to generate nutritional adaptations to obligate hematophagy. This tick harbors a microbiome functionally analogous to those observed in other tick species, dominated by intracellular bacterial symbionts producing the core set of B vitamins (biotin, riboflavin, and folate) essential for tick growth and survival. However, while in other tick species, as in other obligate blood feeders, production of all three core B vitamins is performed by a single nutritional symbiont (Binetruy et al., 2020; Duron et al., 2017; Duron and Gottlieb, 2020), *H. marginatum* harbors two bacterial partners whose core B vitamins biosynthetic capacities depend on a combination of genes, with indigenous and exogenous phylogenetic origins. This multi-partner nutritional symbiosis is stably maintained at least at the tick species level, as supported by our founding of genetically similar FLE and *Midichloria* present in *H. marginatum* populations from geographically distant regions.

Genome reconstructions confirmed the key role of symbionts in *H. marginatum* nutrition, but contrary to other tick species, no single symbiont does compensate for all nutritional needs. The FLE-Hmar genomes contain the genes to synthesize folate and riboflavin but its biotin synthesis pathway is degraded, and thus non-functional. Genome reduction is a conserved trait in all FLEs, but this process has extended in the FLE of *H. marginatum,* with an additional loss of genes in most COG functional categories. This process could have been triggered by the pseudogenization of the mismatch repair system (*mutSL*), leaving the FLE of *H. marginatum* with only one DNA repair system, nucleotide excision repair (*uvrABCD*). Absence of the *mutSL* system, coupled with strict clonality and a small population size during transovarial transmission, have probably led the FLE-Hmar genomes to accumulate deleterious mutations, that resulted in maladaptation, and ultimately limited their beneficial contributions to ticks.

In the context of FLE genome degradation, co-infection with *Midichloria* prevents the breakdown and collapse of nutritional symbiosis in *H. marginatum*; i.e., the *Midichloria* genome contains an intact biotin biosynthesis operon, and thus compensates for genome decay of FLE. Inversely, *Midichloria* has only partial pathways for riboflavin and folate and hence cannot meet the nutritional needs of *H. marginatum* for these B vitamins. Consequently, the co-infection of FLE with *Midichloria* forms a cooperative system essential for the nutritional symbiosis in *H. marginatum*. In addition, complementary heme synthesis genes are present in the FLE and *Midichloria* genomes, and these symbionts may be an additional source of heme, perhaps conferring a fitness advantage to *H. marginatum* during periods of starvation. In animals, heme is essential for the function of all aerobic cells, playing an important role in controlling protein synthesis and cell differentiation. Complete biosynthetic heme gene pathways exist in most animals, but not in ticks which need an exogenous source of heme (Gulia-Nuss et al., 2016; Jia et al., 2020). Vertebrate blood is usually considered to be their unique source for this essential cofactor (Perner et al., 2016), however ticks spend most of their life off host, and thus the presence of heme genes in FLE and *Midichloria* genomes may also indicate symbiont provisioning of heme in *H. marginatum*.

The co-localization of FLE and *Midichloria* in tick ovaries suggests that vertical transmission has led to their stable co-inheritance through tick generations, linking their evolutionary fate, as observed in other symbiotic systems of arthropods (McCutcheon et al., 2019; Vautrin and Vavre, 2009). This co-transmission fidelity creates the conditions under which cooperative interactions can emerge: The stable coexistence of FLE and *Midichloria* genomes makes various genes redundant, as we observed for B vitamin and heme synthesis pathways.

Furthermore, the covariance of FLE and *Midichloria* densities in ticks is positive, suggesting that these two symbionts are not in direct competition but rather cooperative. However, while the nutritional association of FLE with *Hyalomma* species has persisted for millions of years (Azagi et al., 2017), the nutritional role of *Midichloria* seems more recent in this tick genus. Co-infection with *Midichloria* was reported in at least nine *Hyalomma* species (cf. Table S4), but in most species, such as *H. excavatum* and *H. impeltatum* (Azagi et al., 2017; Selmi et al., 2019)*, Midichloria* is present at much lower frequencies than would be expected for an obligate nutritional mutualist, suggesting that it is instead a facultative (i.e., not essential) association. Interestingly, the *Hyalomma* species apparently devoid of *Midichloria*, *H. asiaticum*, harbors a FLE with intact biotin, folate, and riboflavin synthesis pathways (Buysse and Duron, 2021). In *H. marginatum*, the decay of the FLE genome may have been accelerated by the presence of *Midichloria* and the redundancy of biotin biosynthesis, a process also reported in sap-feeding insects (Husnik and Mccutcheon, 2016; Manzano-Marín et al., 2020). While it is clear that two events have led to the current status (loss of biotin genes and acquisition of *Midichloria*), the order by which these happened is unclear. *Midichloria* could have rescued a decaying symbiosis, or could have led, with its entrance in the system, to relaxation of selective constraints on FLE. Based on the common presence of *Midichloria* at lower frequencies in other *Hyalomma* species, a scenario of gradual parallel occurrence of these two events cannot be ruled out.

Our analyses also indicated that the biotin operon detected in the *Midichloria* of *H. marginatum* has an exogenous origin, being acquired following an ancient lateral gene transfer pre-dating the divergence between the *Midichloria*of *H. marginatum* and the one of *I. ricinus*. Accumulating genomic sequences indicate that this streamlined biotin operon (denoted in some studies as the Biotin synthesis Operon of Obligate intracellular Microbes – BOOM (Driscoll et al., 2020)) has experienced transfers between distantly related endosymbionts inhabiting arthropod cells including *Wolbachia* (Driscoll et al., 2020; Gerth and Bleidorn, 2016; Ju et al., 2020; Nikoh et al., 2014), *Rickettsia* (Gillespie et al., 2012), *Cardinium* (Penz et al., 2012) and *Legionella* (Říhová et al., 2017). Remarkably, the invasive nature of this streamlined biotin operon has contributed not only to the emergence of novel nutritional symbioses in *H. marginatum* with *Midichloria*, but also in other blood feeders, such as bedbugs with *Wolbachia* (Nikoh et al., 2014) and rodent lice with *Legionella* (Říhová et al., 2017), or in sap feeders as white flies with *Cardinium* (Santos-Garcia et al., 2014) and planthoppers with *Wolbachia* (Ju et al., 2020). Until recently, lateral gene transfer was thought to be rare in intracellular symbionts, as they reside in confined and isolated environments. However, coinfections of unrelated endosymbionts within the same host cell have created freely recombining intracellular bacterial communities, a unique environment previously defined as the ‘intracellular arena’ (Kent and Bordenstein, 2010). The detection of the streamlined biotin operon in *Wolbachia*, *Rickettsia*, and *Cardinium* – three of the most common intracellular symbionts of arthropods (Duron et al., 2008) – corroborates the role of such ‘intracellular arenas’ in the evolutionary dynamics of nutritional symbioses, especially as the biotin operon can be found on plasmids (Gillespie et al., 2012).

The macroevolutionary and ecological consequences of acquiring nutritional symbionts are immense, ticks would simply not exist as strict blood feeders without such mutualistic interactions (Duron and Gottlieb, 2020). Our investigations corroborate that nutritional symbionts of ticks have all converged towards an analogous process of B vitamin provisioning. While past investigations showed that this evolutionary convergence consists in severe degeneration of symbiont genomes accompanied by preservation of biotin, folate, and riboflavin biosynthesis pathways (Buysse and Duron, 2021; Duron and Gottlieb, 2020), nutritional endosymbiosis in *H. marginatum* is influenced by a more complex web of interactions resulting in a dynamic mosaic of symbiont combinations and offering novel evolutionary possibilities.

## Materials and Methods

### Tick collection, DNA extraction and molecular screening

Adult females of *H. marginatum* were collected from domestic animals from three geographical regions (Italy, Spain, and Israel) (Figure 1). Whole body of one female from Italy was processed for metagenome sequencing, while ovaries from females collected in Spain (n=10) and Israel (n=10) were used for metagenome sequencing (and for qPCR in the case of samples from Spain). For each sample, total DNA was extracted using either the DNeasy Blood and Tissue kit (QIAGEN) or the NucleoSpin Plant II kit (Macherey-Nagel).

A molecular screening was performed by qPCR (iQ5 system, Bio-Rad) on Spanish samples to evaluate the bacterial load of FLE and *Midichloria* symbionts in ovaries. For each specimen, three SYBR Green-based qPCR (Bio-Rad) assays were performed: (i) A specific amplification for the *H. marginatum cal* gene, (ii) a specific amplification for the FLE *rpoB* gene, and (iii) a specific amplification for the *Midichloria gyrB* gene. Thermal protocols and primers are listed in Supplementary Table S5.

### Genome sequencing, assembly and annotation

Metagenomes were sequenced on Illumina HiSeq 2500, after a library preparation performed with Nextera XT. For the Italian and the Spanish samples, the raw paired-end reads quality was evaluated with FastQC (Andrews, 2010). Paired-end reads were trimmed via Trimmomatic (v0.40) (Bolger et al., 2014) and then assembled using SPAdes (v3.10.0) (Bankevich et al., 2012) following the Blobology pipeline (Kumar et al., 2013) (thus separating reads according to G+C content, coverage, and taxonomic annotation). Discarding of contaminating sequences and refining of the assemblies were performed manually through the analysis of the assemblies’ graphs with Bandage (v0.8.1) (Wick et al., 2015). Bacterial reads were separated from the tick reads, allowing the assembling of different bacterial genomes (“filtered” assemblies). For the Israeli sample, reads were trimmed using Cutadapt (v1.13) (Martin, 2011) and contaminants’ reads were removed through a mapping against Coliphage phi-X174 (GenBank: NC_001422) and *Staphylococcus aureus* (NC_010079) genomes with BWA (v0.7.17-r1188) (option mem) and SAMtools (v1.10) (Li and Durbin, 2009). Remaining reads were further assembled using SPAdes (v3.11.1) (Bankevich et al., 2012) with default parameters. These “raw” assembled scaffolds were compared to different reference genomes separately (FLE of *O. moubata* FLEOm, *M. mitochondrii*, and *R. aeschlimannii*; Table S2) to discard unspecific scaffolds using RaGOO (v1.02) (Alonge et al., 2019), and, for each reference genome separately, the assignation of the remaining scaffolds (“filtered” assembly) was confirmed using the online NCBI BLAST tool (https://blast.ncbi.nlm.nih.gov/Blast.cgi). The quality assessment and the completeness of the “filtered” assemblies from all *H. marginatum* metagenomes were estimated using QUAST (v4.6.3) (Gurevich et al., 2013) and miComplete (v1.1.1) (Hugoson et al., 2020) with the -- hmms Bact105 setting. The newly obtained genomes were annotated using Prokka (v1.13.1) (Seemann, 2014) with default parameters. The new *Midichloria* genomes were annotated using a manually revised annotation of *M. mitochondrii* (Table S2) as a reference.

### Genomic content, structure, and metabolic pathways analyses

The FLE and *Midichloria* whole-genome alignments were obtained separately using Mauve (Darling et al., 2004). Previously, contigs from newly obtained genomes were reordered with Mauve. Graphical representations of newly sequenced genomes were produced using CGView (v1.5) (Stothard and Wishart, 2005). Genes involved in biosynthesis pathways (B vitamins and heme), *Francisella* Pathogenecity Island (FPI), and mismatch repair systems (*mutSL* and *uvrABCD*) were retrieved using BLASTn, BLASTp, and tBLASTx on both newly obtained and reference FLE and *Midichloria* genomes. Pseudogenes prediction was performed using Pseudofinder (version downloaded in date April 1^st^ 2020) (Syberg-Olsen et al., 2020) on each genome, including newly obtained and reference genomes of FLE and *Midichloria* (Table S2). The FLE and *Midichloria* datasets were filtered excluding pseudogenes from the subsequent analyses. Clusters of orthologous groups of proteins (COGs) were predicted using the NCBI pipeline (Galperin et al., 2015) and COGs presence was then plotted with an in-house R script. Then, two different datasets were created: (i) a FLE dataset including the three newly sequenced FLE-Hmar genomes and other published FLE genomes; and (ii) a *Midichloria* dataset including the three newly sequenced Midi-Hmar genomes and *M. mitochondrii*. For each dataset, single-copy orthologs (SCO) were identified using OrthoFinder (v2.3.12) (Emms and Kelly, 2019), then SCO lists were retrieved on a R environment (v3.6.2) to build Venn diagrams using the “VennDiagram” R package (Chen and Boutros, 2011).

### Phylogenomics

The FLE, *Midichloria* and *Rickettsia* datasets were filtered excluding the pseudogene sequences and were then used for the phylogenetic analyses. For each symbiont dataset, single-copy orthologs (SCO) were identified using OrthoFinder (v2.3.12) (Emms and Kelly, 2019) and then aligned with MUSCLE (v3.8.31) (Edgar, 2004). Poorly aligned positions and divergent regions were removed by Gblocks (v0.91b) (Castresana, 2000) prior to concatenation of individual alignments using an in-house script. Finally, the most suitable evolutionary model was determined using modeltest-ng (v0.1.5) (Darriba et al., 2020) according to the Akaike information criterion and maximum likelihood (ML) trees were inferred using RAxML (v8.2.9) (Stamatakis, 2014) with 1,000 bootstrap replicates.

### Biotin operon phylogeny and structure

Nucleic and protein sequences of the six genes of the biotin biosynthesis pathway were retrieved using BLASTn, BLASTp, and tBLASTx on all *Midichloria* genomes. Sequences from described streamlined biotin operons and sequences from biotin genes from other symbionts (retrieved from GenBank) were included in the dataset (details in Table S3). All protein sequences of the six biotin genes were manually concatenated and aligned with MAFFT (v7.450) (Katoh et al., 2002) and ambiguous positions were removed using trimAl (v1.2rev59) (Capella-Gutiérrez et al., 2009). The appropriate substitution model, and then ML-tree, were determined as described above. Meanwhile, 16S rRNA sequences from all organisms included in this dataset were retrieved from GenBank and aligned with MEGA software. The Gblocks program with default parameters was used to obtain non-ambiguous sequence alignments. Phylogenetic analyses were performed with MEGA software, i.e. determination of the appropriate substitution model using Akaike information criterion, ML-tree inference. Clade robustness was assessed by bootstrap analysis using 1,000 replicates.

Contigs carrying the streamlined biotin operon were retrieved from newly obtained and reference genomes of different intracellular bacteria. Contigs were annotated using Prokka (v1.13.1) (Seemann, 2014) with default parameters. Prokka GenBank outputs were imported in R (v3.6.2) and analyzed using the “genoplotR” R package (Guy et al., 2010). Sequences of *ribH* and *folD* genes from bacteria carrying the streamlined biotin operon and from other bacteria were secondarily separately analyzed. Phylogenetic analyses on these two genes were performed as described above for biotin genes.

### Bibliographic survey and phylogeny of *Hyalomma*

To access the associations between *Hyalomma* species with FLE and *Midichloria*, we performed an extensive bibliographic survey of published studies. For *Hyalomma* species for which presence/absence of FLE and *Midichloria* is documented, we retrieved cytochrome c oxidase I (*cox1*) gene sequences from GenBank and further performed a ML phylogenetic analysis with MEGA software as described above.

### Statistical analyses

Statistical analyses were carried out using the R statistical package (v3.6.2) and the “lme4” R package (Bates et al., 2015). The FLE load was analyzed using linear mixed-effects models on a dataset containing loads measured in ovaries from adult engorged ticks. The normality of the data distribution was tested through a Shapiro-Wilk test with a 95% confidence. Outliers were removed from datasets. To build models, *Midichloria* load was fitted as a fixed explanatory variable. The best-fitted model was determined with the following procedure: the maximal model, including all higher-order interactions, was simplified by sequentially eliminating interactions and non-significant terms to establish a minimal model. A likelihood ratio test (LRT) (using a chi-square distribution and a p-value cut-off of 0.05) was performed to establish the difference between sequential models. The comparison between the densities of FLE and *Midichloria* of *H. marginatum* was tested by a paired t test.

## Data and code availability

The bacterial genomes obtained in this study were deposited in GenBank under accession numbers GCA_910592745 (FLE-Hmar-ES), GCA_910592815 (FLE-Hmar-IL), GCA_910592705 (FLE-Hmar-IT), GCA_910592695 (Midi-Hmar-ES), GCA_910592725 (Midi-Hmar-IL), GCA_910592865 (Midi-Hmar-IT) and GCA_910592765 (RAES-Hmar-IT). Command lines used for the different genomic analyses are available on GitHub (https://github.com/mariebuysse/Hmar-2021).

## Acknowledgements

We are grateful to Tal Azagi for tick collection and extraction and Inbar Plaschkes for initial sequence analysis of the Israeli sample, to Justine Grillet, Valentina Serra and Catherine Hartley for technical support during molecular analyses. This work benefited from (1) an international ‘Joint Research Projects’ grant (EVOSYM) co-managed by the Ministry of Science, Technology and Space (Israel) and the Centre National de la Recherche Scientifique (CNRS, France), from (2) The Israel Science Foundation (ISF grant No. 1074/18), and from (3) the Human Frontier Science Programme Grant RGY0075/2017 to DS, and the Italian Ministry of Education, University and Research (MIUR): Dipartimenti di Eccellenza Programme (2018–2022) – Department of Biology and Biotechnology “L. Spallanzani”, University of Pavia to DS.

## Author contributions

D.S., Y.G. and O.D. designed the study. M.B., A.F., Y.G., D.S. and O.D. wrote the manuscript with input from other authors. M.B., A.F., T.N. and F.C. performed the genomic, phylogenomic and statistical analyses. F.C., E.O., A.G, A.P., B.M., C.B. and A.C. participated to collection and identification of specimen collection, or molecular biology. All authors agreed on the final version of the manuscript.

## SUPPLEMENTARY DATA

### Supplementary table

**Table S1.** Summary of the assembly information and quality analyses of the newly sequenced genomes of FLE, *Midichloria* and *Rickettsia* bacteria. Only complete genomes are shown and the partial genome of *R. aeaschlimannii* (RAES-Hmar-IS) was not presented.

| Specimen details |  |  | Assembling information |  |  |  |  |  |  |  |  |
| --- | --- | --- | --- | --- | --- | --- | --- | --- | --- | --- | --- |
| Genome | Isolate | Origin | Genome size (bp) | Contigs number | N50 | L50 | Completeness (%) | GC rate (%) | Genes | Pseudogenes | Coding density (%) |
| *Francisella*-LE of *Hyalomma marginatum* | FLE-Hmar-ES | Spain | 1510123 | 17 | 254696 | 3 | 0,9905 | 31,11 | 937 | 795 | 54 |
| *Francisella*-LE of *Hyalomma marginatum* | FLE-Hmar-IT | Italy | 1510130 | 13 | 382079 | 2 | 0,9905 | 31,11 | 934 | 789 | 54 |
| *Francisella*-LE of *Hyalomma marginatum* | FLE-Hmar-IL | Israel | 1536710 | 25 | 106485 | 5 | 0,9905 | 31,23 | 961 | 798 | 54 |
| *Midichloria* of *Hyalomma marginatum* | Midi-Hmar-ES | Spain | 1168604 | 179 | 19210 | 20 | 0,981 | 35,03 | 1071 | 192 | 78 |
| *Midichloria* of *Hyalomma marginatum* | Midi-Hmar-IT | Italy | 1130547 | 359 | 10359 | 30 | 0,9524 | 35,23 | 1032 | 192 | 76 |
| *Midichloria* of *Hyalomma marginatum* | Midi-Hmar-IL | Israel | 1144281 | 123 | 12627 | 30 | 0,981 | 35,03 | 1055 | 190 | 79 |
| *Rickettsia aeschlimannii* (of *H. marginatum*) | RAES-Hmar-IT | Italy | 1356794 | 165 | 67640 | 4 | 0,9429 | 32,32 | 1122 | 475 | 68 |

**Table S2.**
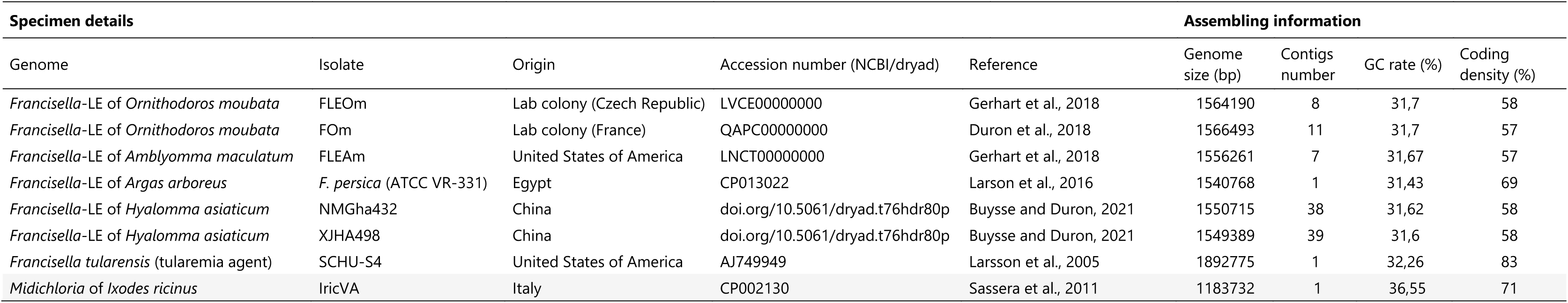
Summary of the information on the NCBI reference genomes employed for the characterization of the FLE and *Midichloria* genomes.

**Table S3.**
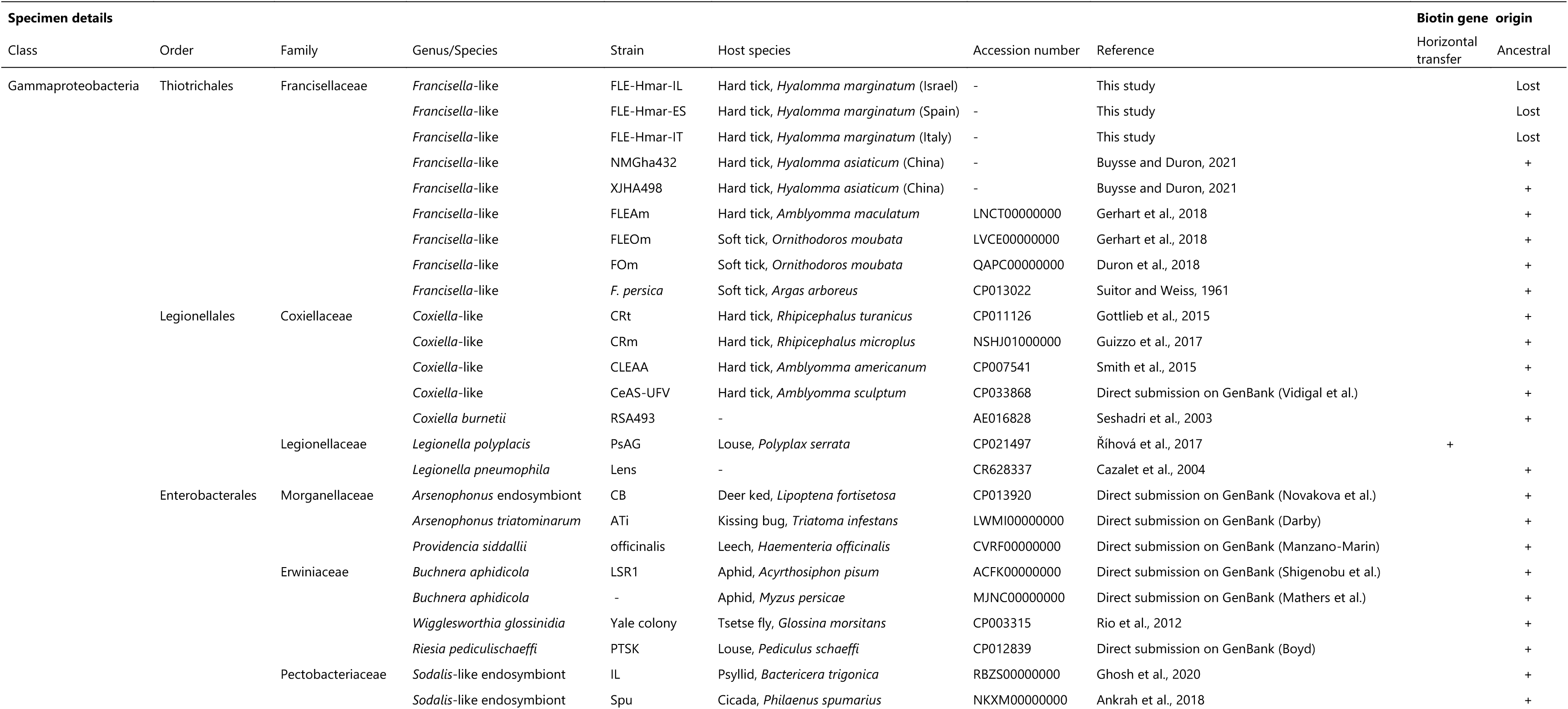
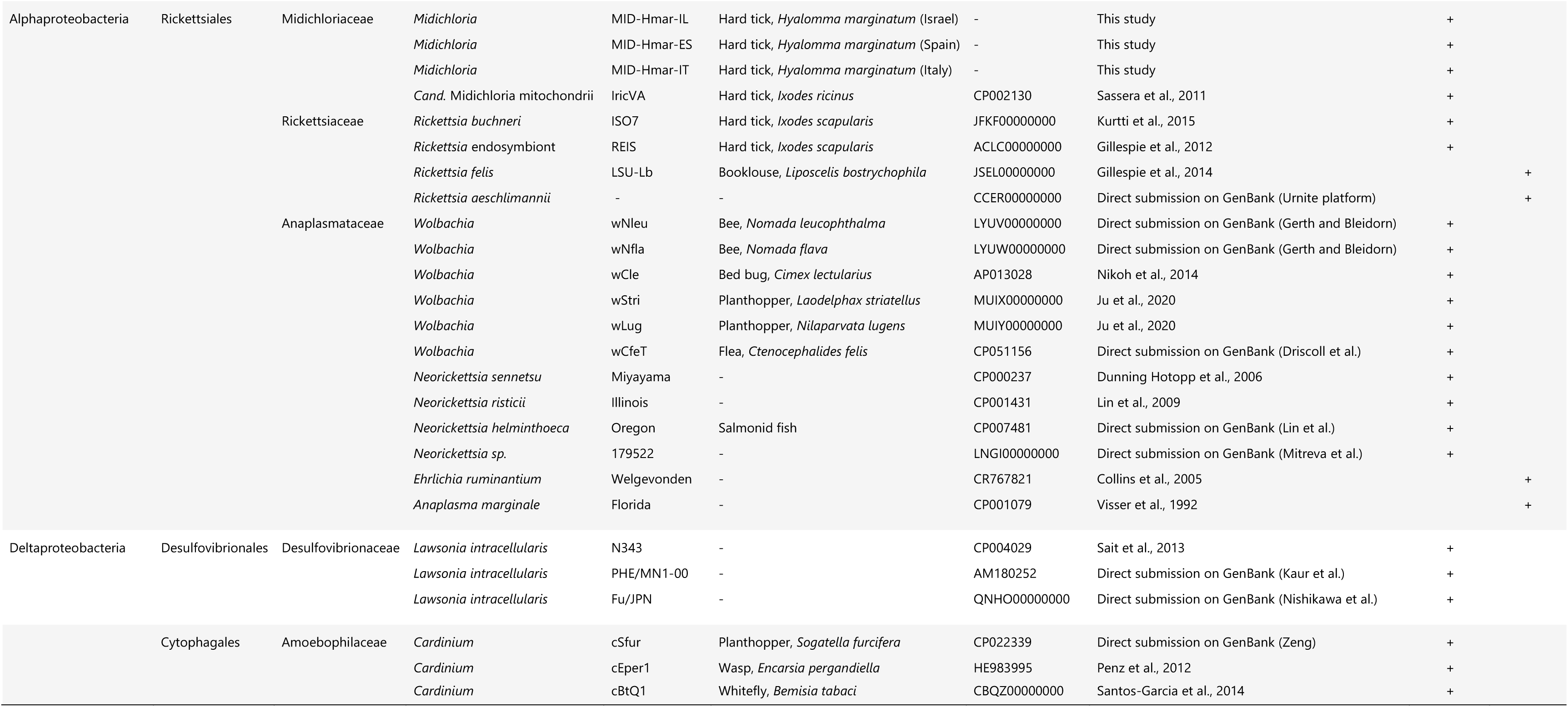
Details regarding symbionts used in phylogenetic analyses of B vitamin biosynthetic pathways and streamlined biotin operon of FLE and *Midichloria*.

**Table S4.** Illustrative (non-exhaustive) studies investigating the presence of FLE and *Midichloria* symbionts across *Hyalomma* tick species. +: presence; –: absence; ND: not determined.

| Specimen details |  | Detection details |  |  | Reference |
| --- | --- | --- | --- | --- | --- |
| Tick species | Geographic origin | Symbiont presence |  | Detection method |  |
|  |  | FLE | *Midichloria* |  |  |
| *Hyalomma* | Israel | + | - | PCR | Azagi et al., 2017 |
| *aegyptium* | Anatolia, Mugla Province | + | ND | Metagenomic analysis | Brinkmann et al., 2019 |
|  | Qatar, Doha | ND | + | PCR | Barradas et al., 2020 |
|  | Bulgaria | + | ND | PCR | Ivanov et al., 2011 |
| *H. anatolicum* | Turkey | - | - | PCR | Duron et al., 2017 |
|  | Turkey | ND | + | PCR & qPCR | Cafiso et al., 2016 |
|  | Pakistan, Punjab Province | + | + | Bacterial barcoding | Adegoke et al., 2020 |
| *H. asiaticum* | China, Xinjiang Province | + | ND | PCR | Wang et al., 2018 |
|  | China | + | - | Metagenomic analysis | Buyse and Duron, 2021 |
| *H. dromedarii* | Israel | + | ND | PCR | Azagi et al., 2017 |
|  | Saudi Arabia | + | ND | PCR | Elbir et al., 2020 |
|  | Tunisia, Sousse | ND | + | PCR | Selmi et al., 2019 |
|  | Tunisia, Kebili | ND | - | PCR | Selmi et al., 2019 |
|  | Egypt | + | ND |  | Ghoneim et al., 2017 |
|  | United Arab Emirates, Al-Ain | + | ND | Bacterial barcoding | Perveen et al., 2020 |
|  | Palestine, Jericho | + | - | Metagenomic analysis | Ravi et al., 2018 |
|  | Saudi Arabia, Al Hofuf | + | - | PCR & bacterial barcoding | Elbir et al., 2019 |
|  | Saudi Arabia, Hail Province | + | - | Bacterial barcoding | Alreshidi et al., 2020 |
| *H. excavatum* | Israel | + | ND | PCR | Azagi et al., 2017 |
|  | Tunisia, Kebili | ND | - | PCR | Selmi et al., 2019 |
|  | Turkey, Mecitözü | + | + | PCR | Duron et al., 2017 |
|  | Turkey | ND | + | PCR & qPCR | Cafiso et al., 2016 |
|  | Egypt | ND | + | Reverse line blotting (RLB) | AL-Hosary et al., 2021 |
| *H. impeltatum* | Tunisia, Gabes | ND | + | PCR | Selmi et al., 2019 |
|  | Tunisia, Sousse | ND | + | PCR | Selmi et al., 2019 |
|  | Tunisia, Kebili | ND | - | PCR | Selmi et al., 2019 |
|  | Tunisia, Kairouan | ND | - | PCR | Selmi et al., 2019 |
|  | Tunisia, Tataouine | ND | - | PCR | Selmi et al., 2019 |
|  | Zimbabwe | + | - | PCR | Duron et al., 2017 |
| *H. lusitanicum* | Spain | + | + | Bacterial barcoding | Díaz-Sánchez et al., 2021 |
|  | Canaries, Veneguera | + | - | PCR | Duron et al., 2017 |
|  | Spain, Madrid | + | ND | PCR | Lopes de Carvalho et al., 2016 |
|  | Portugal | - | ND | PCR | Lopes de Carvalho et al., 2016 |
| *H. marginatum* | Israel | + | ND | PCR & FISH | Azagi et al., 2017 |
|  | Anatolia, Mugla Province | + | ND | Illumina HiSeq sequencing | Brinkmann et al., 2019 |
|  | France | + | + | PCR | Duron et al., 2017 |
|  | Italy, Sicilia | ND | + | PCR | Epis et al., 2008 |
|  | Ventotene Island (Mediterranean Sea) | ND | + | PCR | Di Lecce et al., 2018 |
|  | Bulgaria | + | ND | PCR | Ivanov et al., 2011 |
|  | Spain, Madrid | - | ND | PCR | Lopes de Carvalho et al., 2016 |
|  | Portugal | - | ND | PCR | Lopes de Carvalho et al., 2016 |
| *H. rufipes* | Israel | + | ND | PCR | Azagi et al., 2017 |
|  | China, Gansu Province | - | + | High-throughput sequencing of small RNAs | Luo et al., 2017 |
|  | Ethiopia, Didessa Valley | + | ND | PCR | Szigeti et al., 2014 |
| *H. truncatum* | Namibia | + | ND | PCR | Direct submission on GenBank (JF290387) |
|  | Italy, Sicilia | ND | + | PCR | Epis et al., 2008 |

**Table S5.**
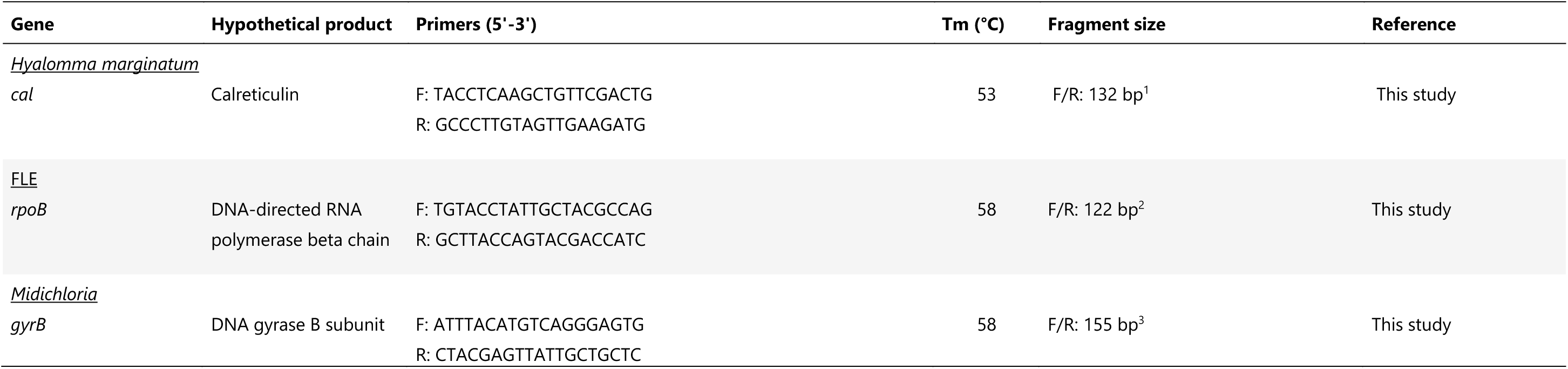
Genes and primers used in this study. Primers were used for qPCR assays conducted for relative quantification of FLE and *Midichloria*. ^1^ PCR conditions: primers concentration 150 nM; thermal profile: 95 °C for 3 min, and 40 cycles at 95 °C for 15 sec, 55 °C for 20 sec, 72 °C for 10 sec; ^2^ PCR conditions: primers concentration 250 nM; thermal profile: 95 °C for 3 min, and 40 cycles at 95 °C for 15 sec, 58 °C for 30 sec; ^3^ PCR conditions: primers concentration 250 nM; thermal profile: 95 °C for 3 min, and 40 cycles at 95 °C for 15 sec, 58°C for 30 sec.

### Supplementary figures

**Figure S1.**
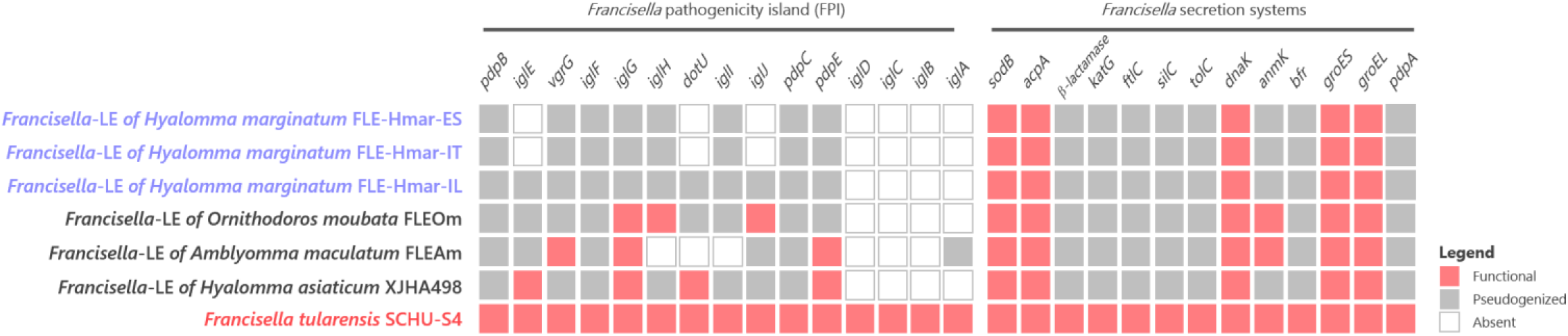
Conservation level of the genes involved in the *Francisella* pathogenicity island (FPI) on FLE of *Hyalomma marginatum* and references genomes. Genomes of FLE of *Hyalomma marginatum* are indicated by a blue font. *Francisella* pathogenic species is indicated by a red font. Red squares, functional genes; grey squares, pseudogenes; white squares, missing genes.

**Figure S2.**
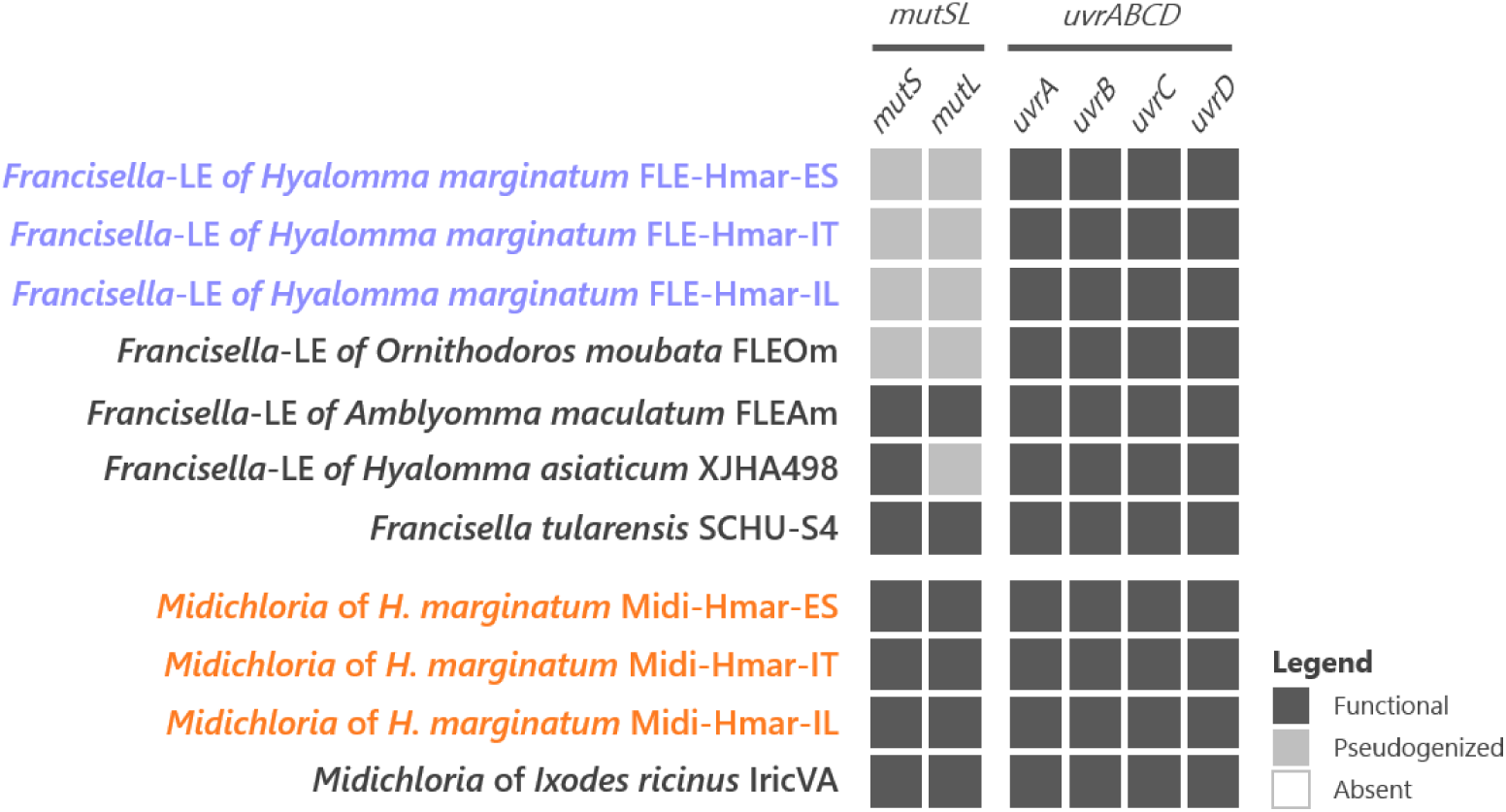
Conservation level of the genes involved in the mismatch repair system (*mutSL*) and the nucleotide excision repair system (*uvrABCD*) on FLE and *Midichloria* of *Hyalomma marginatum* and references genomes. Newly sequenced FLE and *Midichloria* genomes are indicated by a blue and an orange font, respectively. Black squares, functional genes; grey squares, pseudogenes; white squares, missing genes.

**Figure S3.**
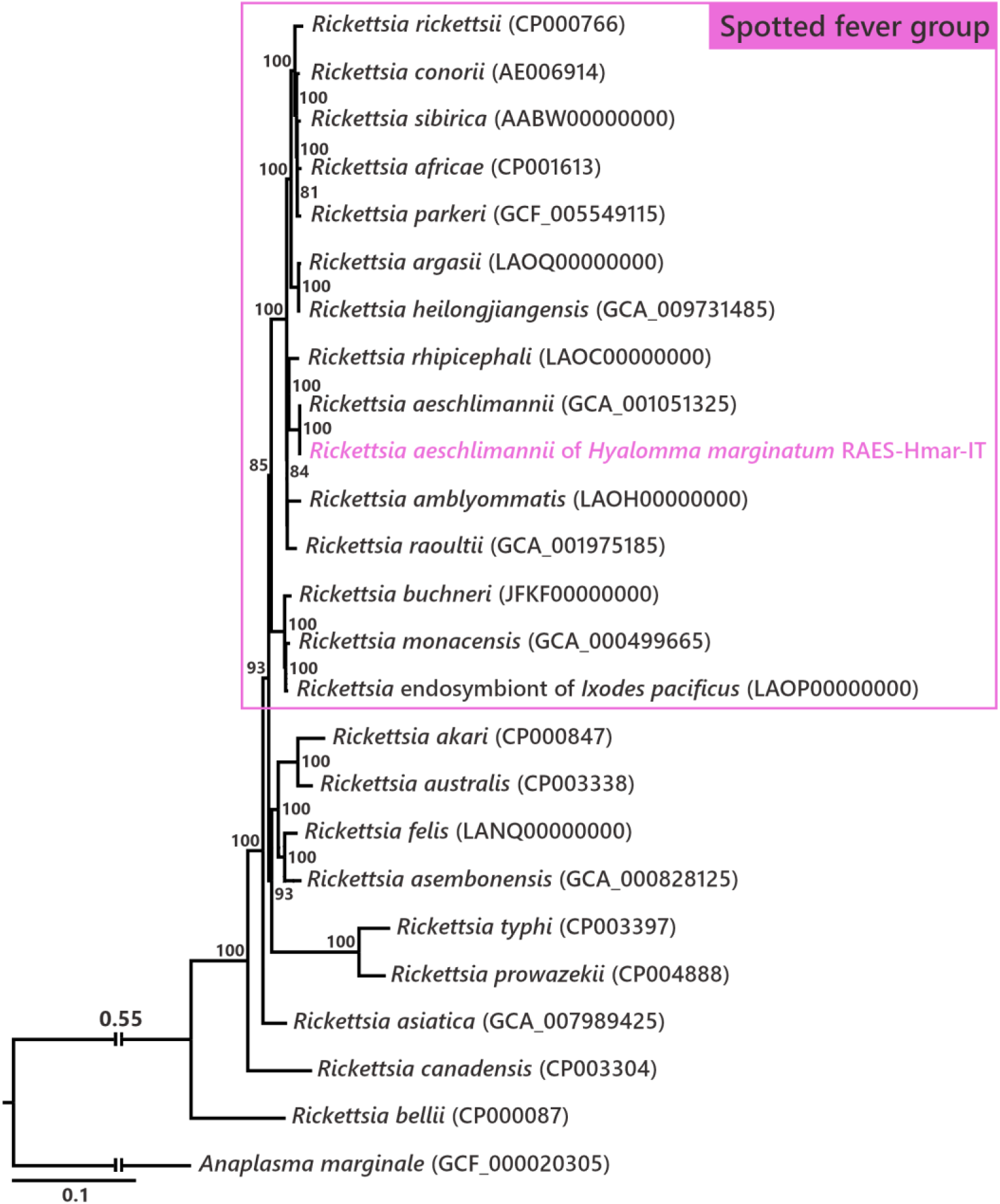
*Rickettsia* phylogenetic tree assessing the affiliation of the *Rickettsia* detected in *Hyalomma marginatum* specimens from Italy and Israel. The phylogenetic tree was inferred using maximum likelihood from a concatenated alignment of 324 single-copy orthologous genes (88,128 amino acids; best-fit approximation for the evolutionary model: JTT+G4+I). The numbers on each node represent the support of 1,000 bootstrap replicates; only bootstrap values >70% are shown. The scale bar is in units of substitution/site. Sequence from the newly sequenced *Rickettsia* genome is indicated by a pink font.

**Figure S4.**
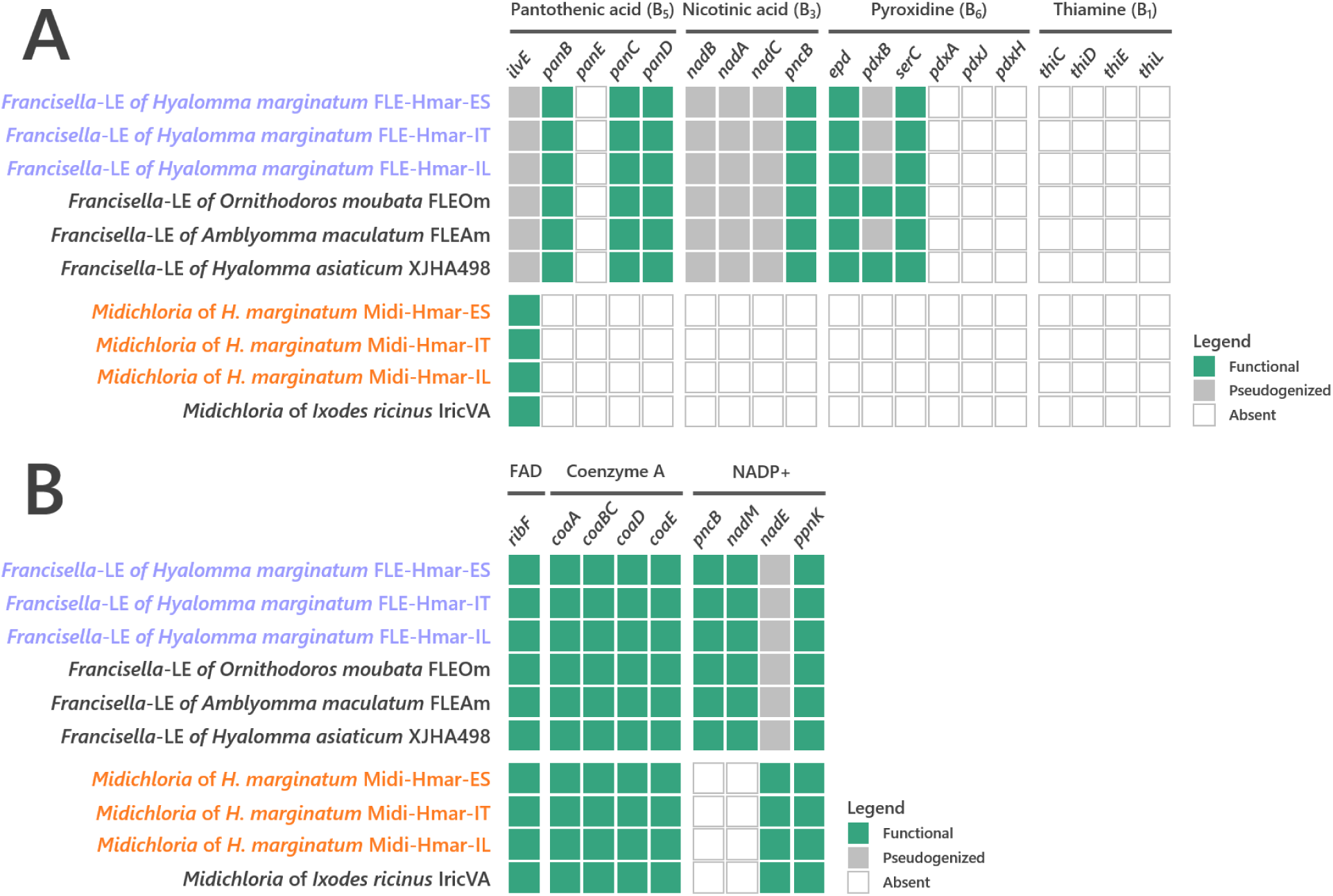
Conservation level of (A) pantothenic acid, nicotinic acid, pyroxidine, and thiamine B vitamins and (B) cofactors biosynthetic pathways of FLE and *Midichloria* symbionts. Newly sequenced FLE and *Midichloria* genomes are indicated by a blue and an orange font, respectively. Green squares, functional genes; grey squares, pseudogenes; white squares, missing genes.

**Figure S5.**
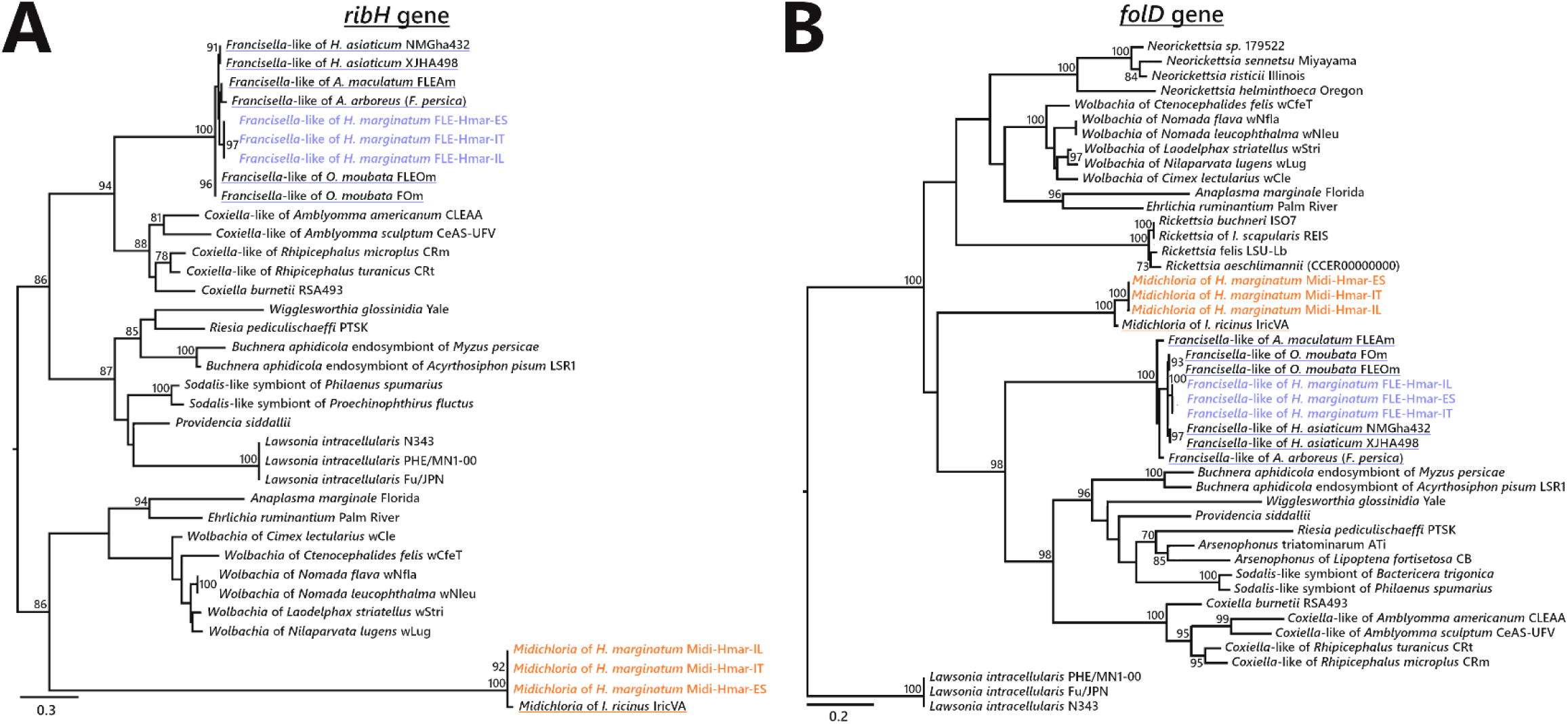
Phylogenetic trees of representative genes of riboflavin and folate biosynthesis pathways of FLE and *Midichloria*. The phylogenetic trees of (A) *ribH* (riboflavin pathway), and (B) *folD* (folate) were inferred using maximum likelihood from amino acid alignments. The numbers on each node represent the support of 1,000 bootstrap replicates; only bootstrap values >70% are shown. The scale bar is in units of substitution/site. *ribH*: 129 amino acids unambiguously aligned, best-fit approximation for the evolutionary model LG+G4; *folD*: 273 amino acids unambiguously aligned, best-fit approximation for the evolutionary model CPREV+G4+I. Newly sequenced FLE and *Midichloria* sequences are indicated by a blue and an orange font, respectively, while sequences from other FLE and *Midichloria* are highlighted in blue and orange, respectively.

**Figure S6.**
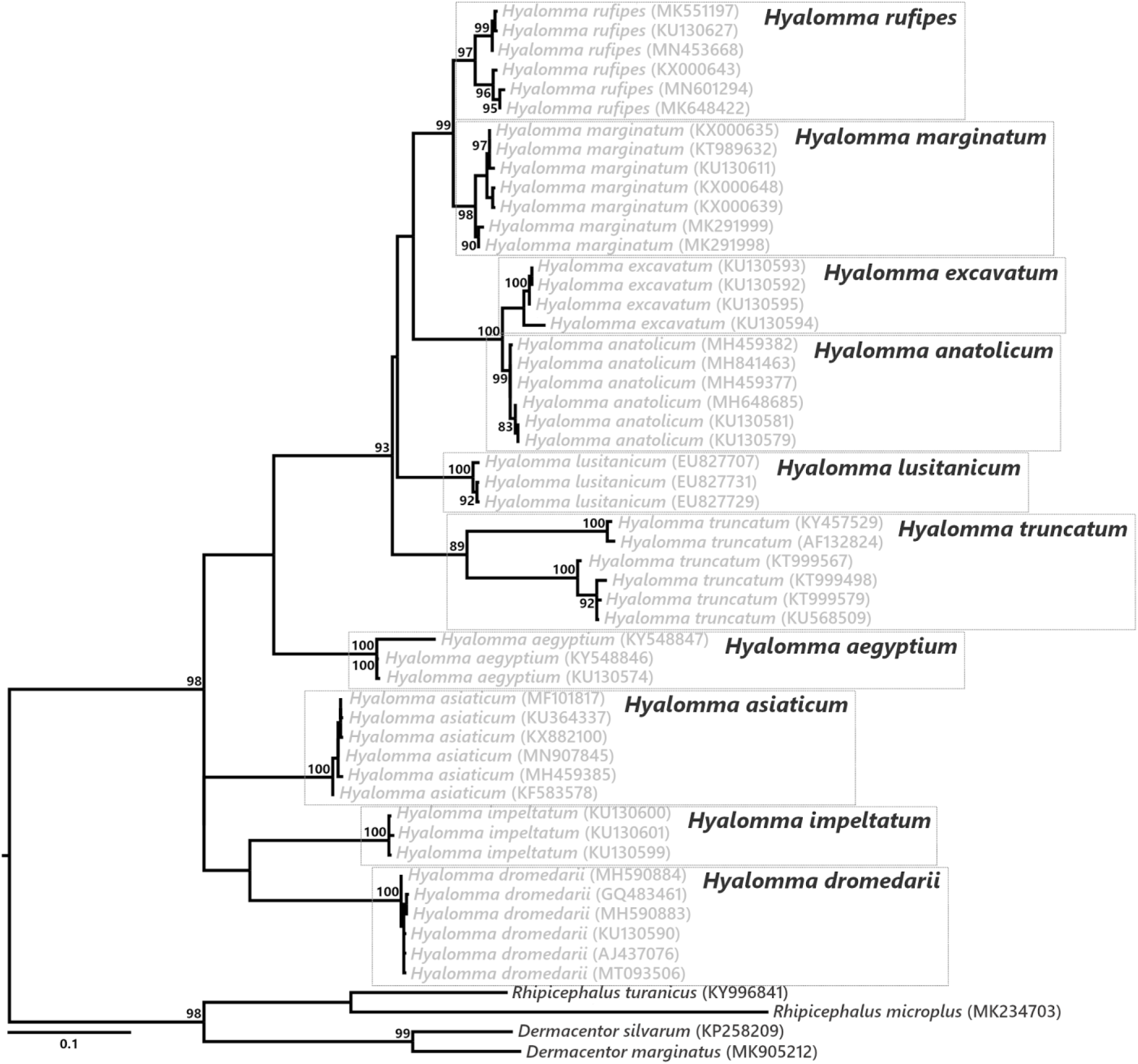
Phylogenetic relationship of the *Hyalomma* species used in our study. The phylogenetic tree was inferred using maximum likelihood from an alignment of the cytochrome c oxidase I (*cox1*) gene sequences (546 bp unambiguously aligned, best-fit approximation for the evolutionary model: HKY+G+I). The numbers on each node represent the support of 1,000 bootstrap replicates; only bootstrap values >80% are shown. The scale bar is in units of substitution/site.

**Figure S7.**
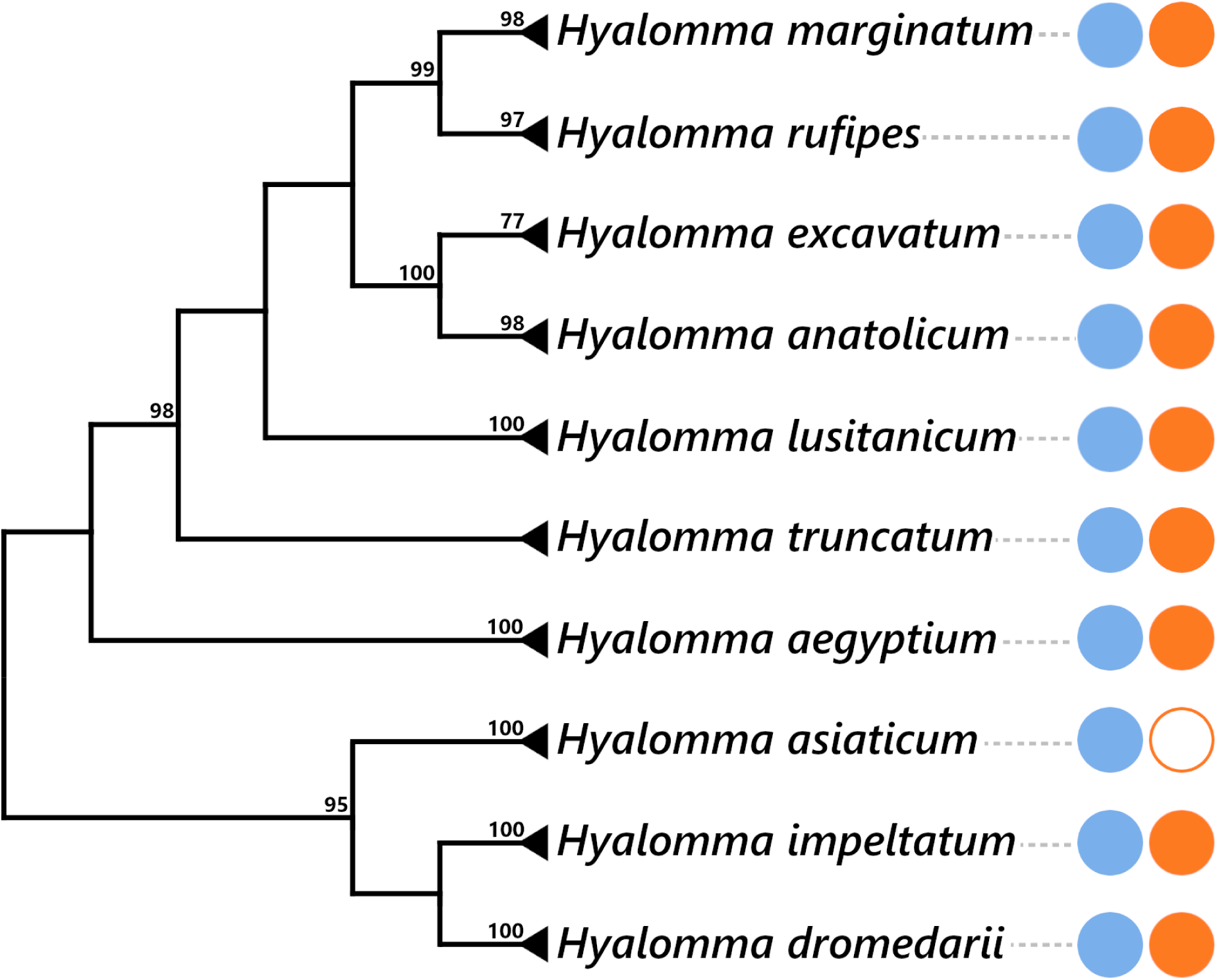
Presence of FLE and *Midichloria* within the *Hyalomma* genus. The *Hyalomma* cladogram was adapted from *cox1* phylogeny showed at Figure S6. The numbers on each node represent the support of 1,000 bootstrap replicates; only bootstrap values >80% are shown. The scale bar is in units of substitution/site. Blue and orange plain circles indicate the presence of FLE and *Midichloria*, respectively, as detailed in Table S4. Blank circle represents an absence of symbiont.

## References

Alonge M, Soyk S, Ramakrishnan S, Wang X, Goodwin S, Sedlazeck FJ, Lippman ZB, Schatz MC. 2019. RaGOO: Fast and accurate reference-guided scaffolding of draft genomes. Genome Biol 20:1–17. doi:10.1186/s13059-019-1829-6

Andrews S. 2010. FastQC: A quality control tool for high throughput sequence data.

Azagi T, Klement E, Perlman G, Lustig Y, Mumcuoglu KY, Apanaskevich DA, Gottlieb Y. 2017. *Francisella*-like endosymbionts and *Rickettsia* species in local and imported *Hyalomma* ticks. Appl Environ Microbiol 83:1–14. doi:10.1128/AEM.01302-17

Bankevich A, Nurk S, Antipov D, Gurevich AA, Dvorkin M, Kulikov AS, Lesin VM, Nikolenko SI, Pham S, Prjibelski AD, Pyshkin A V., Sirotkin A V., Vyahhi N, Tesler G, Alekseyev MA, Pevzner PA. 2012. SPAdes: A new genome assembly algorithm and its applications to single-cell sequencing. J Comput Biol 19:455–477. doi:10.1089/cmb.2012.0021

Bates D, Mächler M, Bolker BM, Walker SC. 2015. Fitting linear mixed-effects models using lme4. J Stat Softw 67. doi:10.18637/jss.v067.i01

Ben-Yosef M, Rot A, Mahagna M, Kapri E, Behar A, Gottlieb Y. 2020. *Coxiella*-Like endosymbiont of *Rhipicephalus sanguineus* is required for physiological processes during ontogeny. Front Microbiol 11:1–16. doi:10.3389/fmicb.2020.00493

Bennett GM, Moran NA. 2015. Heritable symbiosis : The advantages and perils of an evolutionary rabbit hole. PNAS 112:10169–10176. doi:10.1073/pnas.1421388112

Binetruy F, Buysse M, Lejarre Q, Barosi R, Villa M, Rahola N, Paupy C, Ayala D, Duron O. 2020. Microbial community structure reveals instability of nutritional symbiosis during the evolutionary radiation of *Amblyomma* ticks. Mol Ecol 29:1016–1029. doi:10.1111/mec.15373

Bolger AM, Lohse M, Usadel B. 2014. Trimmomatic: A flexible trimmer for Illumina sequence data. Bioinformatics 30:2114–2120. doi:10.1093/bioinformatics/btu170

Bonnet SI, Pollet T. 2020. Update on the intricate tango between tick microbiomes and tick-borne pathogens. Parasite Immunol 1–12. doi:10.1111/pim.12813

Buysse M, Duron O. 2021. Evidence that microbes identified as tick-borne pathogens are nutritional endosymbionts. Cell.

Campbell MA, Leuven JT Van, Meister RC, Carey KM, Simon C, Mccutcheon JP. 2015. Genome expansion via lineage splitting and genome reduction in the cicada endosymbiont *Hodgkinia*. PNAS 112:10192–10199. doi:10.1073/pnas.1421386112

Capella-Gutiérrez S, Silla-Martínez JM, Gabaldón T. 2009. trimAl: A tool for automated alignment trimming in large-scale phylogenetic analyses. Bioinformatics 25:1972–1973. doi:10.1093/bioinformatics/btp348

Castresana J. 2000. Selection of conserved blocks from multiple alignments for their use in phylogenetic analysis. Mol Biol Evol 17:540–552. doi:10.1093/oxfordjournals.molbev.a026334

Chen H, Boutros PC. 2011. VennDiagram: A package for the generation of highly-customizable Venn and Euler diagrams in R. BMC Bioinformatics 12. doi:10.2307/2689606

Darling ACE, Mau B, Blattner FR, Perna NT. 2004. Mauve: Multiple alignment of conserved genomic sequence with rearrangements. Genome Res 14:1394–1403. doi:10.1101/gr.2289704

Darriba Di, Posada D, Kozlov AM, Stamatakis A, Morel B, Flouri T. 2020. ModelTest-NG: A new and scalable tool for the selection of DNA and protein evolutionary models. Mol Biol Evol 37:291–294. doi:10.1093/molbev/msz189

Di Lecce I, Bazzocchi C, Cecere JG, Epis S, Sassera D, Villani BM, Bazzi G, Negri A, Saino N, Spina F, Bandi C, Rubolini D. 2018. Patterns of *Midichloria* infection in avian-borne African ticks and their trans-Saharan migratory hosts. Parasites and Vectors 11:1–11. doi:10.1186/s13071-018-2669-z

Driscoll TP, Verhoeve VI, Brockway C, Shrewsberry DL, Plumer M, Sevdalis SE, Beckmann JF, Krueger LM, Macaluso KR, Azad AF, Gillespie JJ. 2020. Evolution of *Wolbachia* mutualism and reproductive parasitism: Insight from two novel strains that co-infect cat fleas. PeerJ 8:1–39. doi:10.7717/peerj.10646

Duron O, Binetruy F, Noel V, Cremaschi J, McCoy K, Arnathau C, Plantard O, Goolsby J, Perez De Leon AA, Heylen DJA, Raoul Van Oosten A, Gottlieb Y, Baneth G, Guglielmone AA, Estrada-Pena A, Opara MN, Zenner L, Vavre F, Chevillon C. 2017. Evolutionary changes in symbiont community structure in ticks. Mol Ecol 26:2905–2921. doi:10.1111/mec.14094

Duron O, Bouchon D, Boutin S, Bellamy L, Zhou L, Engelstadter J, Hurst GD. 2008. The diversity of reproductive parasites among arthropods: *Wolbachia* do not walk alone. BMC Biol 6:1–12. doi:10.1186/1741-7007-6-27

Duron O, Gottlieb Y. 2020. Convergence of nutritional symbioses in obligate blood-feeders. Trends Parasitol. doi:10.1016/j.pt.2020.07.007

Duron O, Morel O, Noël V, Buysse M, Binetruy F, Lancelot R, Loire E, Ménard C, Bouchez O, Vavre F, Vial L. 2018. Tick-bacteria mutualism depends on B vitamin synthesis pathways. Curr Biol 28:1–7. doi:10.1016/j.cub.2018.04.038

Edgar RC. 2004. MUSCLE: Multiple sequence alignment with high accuracy and high throughput. Nucleic Acids Res 32:1792–1797. doi:10.1093/nar/gkh340

Emms DM, Kelly S. 2019. OrthoFinder: Phylogenetic orthology inference for comparative genomics. Genome Biol 20:1–14. doi:10.1186/s13059-019-1832-y

Galperin MY, Makarova KS, Wolf YI, Koonin E V. 2015. Expanded microbial genome coverage and improved protein family annotation in the COG database. Nucleic Acids Res 43:D261–D269. doi:10.1093/nar/gku1223

Gerhart JG, Auguste Dutcher H, Brenner AE, Moses AS, Grubhoffer L, Raghavan R. 2018. Multiple acquisitions of pathogen-derived *Francisella* endosymbionts in soft ticks. Genome Biol Evol 10:607–615. doi:10.1093/gbe/evy021

Gerhart JG, Moses AS, Raghavan R. 2016. A *Francisella*-like endosymbiont in the Gulf Coast tick evolved from a mammalian pathogen. Sci Rep 6:1–6. doi:10.1038/srep33670

Gerth M, Bleidorn C. 2016. Comparative genomics provides a timeframe for *Wolbachia* evolution and exposes a recent biotin synthesis operon transfer. Nat Microbiol 2. doi:10.1038/nmicrobiol.2016.241

Gillespie JJ, Joardar V, Williams KP, Driscoll T, Hostetler JB, Nordberg E, Shukla M, Walenz B, Hill CA, Nene VM, Azad AF, Sobral BW, Caler E. 2012. A *Rickettsia* genome overrun by mobile genetic elements provides insight into the acquisition of genes characteristic of an obligate intracellular lifestyle. J Bacteriol 194:376–394. doi:10.1128/JB.06244-11

Gottlieb Y, Lalzar I, Klasson L. 2015. Distinctive genome reduction rates revealed by genomic analyses of two *Coxiella*-like endosymbionts in ticks. Genome Biol Evol 7:1779–1796. doi:10.1093/gbe/evv108

Guizzo MG, Parizi LF, Nunes RD, Schama R, Albano RM, Tirloni L, Oldiges DP, Vieira RP, Oliveira WHC, Leite MDS, Gonzales SA, Farber M, Martins O, Vaz IDS, Oliveira PL. 2017. A *Coxiella* mutualist symbiont is essential to the development of *Rhipicephalus microplus*. Sci Rep 7:1–10. doi:10.1038/s41598-017-17309-x

Gulia-Nuss M, Nuss AB, Meyer JM, Sonenshine DE, Roe RM, Waterhouse RM, Sattelle DB, De La Fuente J, Ribeiro JM, Megy K, Thimmapuram J, Miller JR, Walenz BP, Koren S, Hostetler JB, Thiagarajan M, Joardar VS, Hannick LI, Bidwell S, Hammond MP, Young S, Zeng Q, Abrudan JL, Almeida FC, Ayllón N, Bhide K, Bissinger BW, Bonzon-Kulichenko E, Buckingham SD, Caffrey DR, Caimano MJ, Croset V, Driscoll T, Gilbert D, Gillespie JJ, Giraldo-Calderón GI, Grabowski JM, Jiang D, Khalil SMS, Kim D, Kocan KM, Koči J, Kuhn RJ, Kurtti TJ, Lees K, Lang EG, Kennedy RC, Kwon H, Perera R, Qi Y, Radolf JD, Sakamoto JM, Sánchez-Gracia A, Severo MS, Silverman N, Šimo L, Tojo M, Tornador C, Van Zee JP, Vázquez J, Vieira FG, Villar M, Wespiser AR, Yang Y, Zhu J, Arensburger P, Pietrantonio P V., Barker SC, Shao R, Zdobnov EM, Hauser F, Grimmelikhuijzen CJP, Park Y, Rozas J, Benton R, Pedra JHF, Nelson DR, Unger MF, Tubio JMC, Tu Z, Robertson HM, Shumway M, Sutton G, Wortman JR, Lawson D, Wikel SK, Nene VM, Fraser CM, Collins FH, Birren B, Nelson KE, Caler E, Hill CA. 2016. Genomic insights into the *Ixodes scapularis* tick vector of Lyme disease. Nat Commun 7. doi:10.1038/ncomms10507

Gurevich A, Saveliev V, Vyahhi N, Tesler G. 2013. QUAST: Quality assessment tool for genome assemblies. Bioinformatics 29:1072–1075. doi:10.1093/bioinformatics/btt086

Guy L, Kultima JR, Andersson SGE, Quackenbush J. 2010. GenoPlotR: comparative gene and genome visualization in R. Bioinformatics 27:2334–2335. doi:10.1093/bioinformatics/btq413

Hugoson E, Lam WT, Guy L. 2020. MiComplete: Weighted quality evaluation of assembled microbial genomes. Bioinformatics 36:936–937. doi:10.1093/bioinformatics/btz664

Husnik F, Mccutcheon JP. 2016. Repeated replacement of an intrabacterial symbiont in the tripartite nested mealybug symbiosis. PNAS 5416–5424. doi:10.1101/042267

Jia N, Wang J, Shi W, Du L, Sun Y, Zhan W, Jiang JF, Wang Q, Zhang B, Ji P, Bell-Sakyi L, Cui XM, Yuan TT, Jiang BG, Yang WF, Lam TTY, Chang QC, Ding SJ, Wang XJ, Zhu JG, Ruan XD, Zhao L, Wei J Te, Ye RZ, Que TC, Du CH, Zhou YH, Cheng JX, Dai PF, Guo W Bin, Han XH, Huang EJ, Li LF, Wei W, Gao YC, Liu JZ, Shao HZ, Wang X, Wang CC, Yang TC, Huo QB, Li W, Chen HY, Chen SE, Zhou LG, Ni XB, Tian JH, Sheng Y, Liu T, Pan YS, Xia LY, Li J, Zhao F, Cao WC. 2020. Large-scale comparative analyses of tick genomes elucidate their genetic diversity and vector capacities. Cell 182:1328–1340.e13. doi:10.1016/j.cell.2020.07.023

Ju JF, Bing XL, Zhao DS, Guo Y, Xi Z, Hoffmann AA, Zhang KJ, Huang HJ, Gong JT, Zhang X, Hong XY. 2020. *Wolbachia* supplement biotin and riboflavin to enhance reproduction in planthoppers. ISME J 14:676–687. doi:10.1038/s41396-019-0559-9

Katoh K, Misawa K, Kuma KI, Miyata T. 2002. MAFFT: A novel method for rapid multiple sequence alignment based on fast Fourier transform. Nucleic Acids Res 30:3059–3066. doi:10.1093/nar/gkf436

Kent BN, Bordenstein SR. 2010. Phage WO of *Wolbachia*: Lambda of the endosymbiont world. Trends Microbiol 18:173–181. doi:10.1016/j.tim.2009.12.011

Kumar S, Jones M, Koutsovoulos G, Clarke M, Blaxter M. 2013. Blobology: exploring raw genome data for contaminants, symbionts and parasites using taxon-annotated GC-coverage plots. Front Genet 4:1–12. doi:10.3389/fgene.2013.00237

Li H, Durbin R. 2009. Fast and accurate short read alignment with Burrows-Wheeler transform. Bioinformatics 25:1754–1760. doi:10.1093/bioinformatics/btp324

Łukasik P, Nazario K, Van Leuven JT, Campbell MA, Meyer M, Michalik A, Pessacq P, Simon C, Veloso C, McCutcheon JP. 2017. Multiple origins of interdependent endosymbiotic complexes in a genus of cicadas. Proc Natl Acad Sci U S A 115:E226–E235. doi:10.1073/pnas.1712321115

Machado-Ferreira E, Vizzoni VF, Balsemao-Pires E, Moerbeck L, Gazeta GS, Piesman J, Voloch CM, Soares CAG. 2016. *Coxiella* symbionts are widespread into hard ticks. Parasitol Res 115:4691–4699. doi:10.1007/s00436-016-5230-z

Manzano-Marín A, Coeur d’acier A, Clamens AL, Orvain C, Cruaud C, Barbe V, Jousselin E. 2020. Serial horizontal transfer of vitamin-biosynthetic genes enables the establishment of new nutritional symbionts in aphids’ di-symbiotic systems. ISME J 14:259–273. doi:10.1038/s41396-019-0533-6

Martin M. 2011. Cutadapt removes adapter sequences from high-throughput sequencing reads. EMBnet.journal 17:10–12. doi:doi.org/10.14806/ej.17.1.200

Matsuura Y, Moriyama M, Łukasik P, Vanderpool D, Tanahashi M, Meng XY, McCutcheon JP, Fukatsu T. 2018. Recurrent symbiont recruitment from fungal parasites in cicadas. Proc Natl Acad Sci U S A 115:E5970–E5979. doi:10.1073/pnas.1803245115

McCutcheon JP, Boyd BM, Dale C. 2019. The life of an insect endosymbiont from the cradle to the grave. Curr Biol 29:485–495. doi:10.1016/j.cub.2019.03.032

Narasimhan S, Fikrig E. 2015. Tick microbiome: The force within. Trends Parasitol 31:315–323. doi:10.1016/j.pt.2015.03.010

Nardi T, Olivieri E, Kariuki E, Sassera D, Castelli M. 2021. Sequence of a *Coxiella* endosymbiont of the tick *Amblyomma nuttalli* suggests a pattern of convergent genome reduction in the *Coxiella* genus. Genome Biol Evol 13:1–7. doi:10.1093/gbe/evaa253

Nikoh N, Hosokawa T, Moriyama M, Oshima K, Hattori M, Fukatsu T. 2014. Evolutionary origin of insect-*Wolbachia* nutritional mutualism. Proc Natl Acad Sci U S A 111:10257–10262. doi:10.1073/pnas.1409284111

Penz T, Schmitz-Esser S, Kelly SE, Cass BN, Müller A, Woyke T, Malfatti SA, Hunter MS, Horn M. 2012. Comparative genomics suggests an independent origin of cytoplasmic incompatibility in *Cardinium hertigii*. PLoS Genet 8. doi:10.1371/journal.pgen.1003012

Perner J, Sobotka R, Sima R, Konvickova J, Sojka D, de Oliveira PL, Hajdusek O, Kopacek P. 2016. Acquisition of exogenous haem is essential for tick reproduction. Elife 5:1–20. doi:10.7554/eLife.12318

Říhová J, Novaková E, Husník F, Hypša V. 2017. *Legionella* becoming a mutualist: Adaptive processes shaping the genome of symbiont in the louse *Polyplax serrata*. Genome Biol Evol 9:2946–2957. doi:10.1093/gbe/evx217

Santos-Garcia D, Rollat-Farnier PA, Beitia F, Zchori-Fein E, Vavre F, Mouton L, Moya A, Latorre A, Silva FJ. 2014. The genome of *Cardinium* cBtQ1 provides insights into genome reduction, symbiont motility, and its settlement in *Bemisia tabaci*. Genome Biol Evol 6:1013–1030. doi:10.1093/gbe/evu077

Santos-Garcia Di, Juravel K, Freilich S, Zchori-Fein E, Latorre A, Moya A, Morin S, Silva FJ. 2018. To B or not to B: Comparative genomics suggests *Arsenophonus* as a source of B vitamins in whiteflies. Front Microbiol 9:1–16. doi:10.3389/fmicb.2018.02254

Sassera D, Beninati T, Bandi C, Bouman EAP, Sacchi L, Fabbi M, Lo N. 2006. *Candidatus* Midichloria mitochondrii’, an endosymbiont of the *Ixodes ricinus* with a unique intramitochondrial lifestyle. Int J Syst Evol Microbiol 56:2535–2540. doi:10.1099/ijs.0.64386-0

Sassera D, Lo N, Epis S, D’Auria G, Montagna M, Comandatore F, Horner D, Peretó J, Luciano AM, Franciosi F, Ferri E, Crotti E, Bazzocchi C, Daffonchio D, Sacchi L, Moya A, Latorre A, Bandi C. 2011. Phylogenomic evidence for the presence of a flagellum and *cbb3* oxidase in the free-living mitochondrial ancestor. Mol Biol Evol 28:3285–3296. doi:10.1093/molbev/msr159

Seemann T. 2014. Prokka: Rapid prokaryotic genome annotation. Bioinformatics 30:2068–2069. doi:10.1093/bioinformatics/btu153

Selmi R, Ben Said M, Mamlouk A, Ben Yahia H, Messadi L. 2019. Molecular detection and genetic characterization of the potentially pathogenic *Coxiella burnetii* and the endosymbiotic *Candidatus* Midichloria mitochondrii in ticks infesting camels (*Camelus dromedarius*) from Tunisia. Microb Pathog 136:103655. doi:10.1016/j.micpath.2019.103655

Smith TA, Driscoll T, Gillespie JJ, Raghavan R. 2015. A *Coxiella*-like endosymbiontis a potential vitamin source for the lone star tick. Genome Biol Evol 7:831–838. doi:10.1093/gbe/evv016

Stamatakis A. 2014. RAxML version 8: A tool for phylogenetic analysis and post-analysis of large phylogenies. Bioinformatics 30:1312–1313. doi:10.1093/bioinformatics/btu033

Stothard P, Wishart DS. 2005. Circular genome visualization and exploration using CGView. Bioinformatics 21:537–539. doi:10.1093/bioinformatics/bti054

Sudakaran S, Kost C, Kaltenpoth M. 2017. Symbiont acquisition and replacement as a source of ecological innovation. Trends Microbiol. doi:10.1016/j.tim.2017.02.014

Syberg-Olsen M, Garber A, Keeling P, McCutcheon J, Husnik F. 2020. Pseudofinder.

Takeshita K, Yamada T, Kawahara Y, Narihiro T, Ito M, Kamagata Y, Shinzato N. 2019. Tripartite symbiosis of an anaerobic scuticociliate with two hydrogenosome-associtaed endosymbionts, a *Holospora*-related alphaproteobacterium and a methanogenic archaeon. Appl Environ Microbiol 7:1–14.

Vautrin E, Vavre F. 2009. Interactions between vertically transmitted symbionts: cooperation or conflict? Trends Microbiol 17:95–99. doi:10.1016/j.tim.2008.12.002

Vial L, Stachurski F, Leblond A, Huber K, Vourc’h G, René-Martellet M, Desjardins I, Balança G, Grosbois V, Pradier S, Gély M, Appelgren A, Estrada-Peña A. 2016. Strong evidence for the presence of the tick *Hyalomma marginatum* Koch, 1844 in southern continental France. Ticks Tick Borne Dis 7:1162–1167. doi:10.1016/j.ttbdis.2016.08.002

Wick RR, Schultz MB, Zobel J, Holt KE. 2015. Bandage: Interactive visualization of *de novo* genome assemblies. Bioinformatics 31:3350–3352. doi:10.1093/bioinformatics/btv383

Zhong J, Jasinskas A, Barbour AG. 2007. Antibiotic treatment of the tick vector *Amblyomma americanum* reduced reproductive fitness. PLoS One 2:1–7. doi:10.1371/journal.pone.0000405

## Supplementary references

Adegoke A, Kumar D, Bobo C, Rashid MI, Durrani AZ, Sajid MS, Karim S. 2020. Tick-borne pathogens shape the native microbiome within tick vectors. Microorganisms 8:1–16. doi:10.3390/microorganisms8091299

AL-Hosary A, Răileanu C, Tauchmann O, Fischer S, Nijhof AM, Silaghi C. 2021. Tick species identification and molecular detection of tick-borne pathogens in blood and ticks collected from cattle in Egypt. Ticks Tick Borne Dis 12. doi:10.1016/j.ttbdis.2021.101676

Alreshidi MM, Veettil VN, Noumi E, Del Campo R, Snoussi M. 2020. Description of microbial diversity associated with ticks *Hyalomma dromedarii* (Acari: Ixodidae) isolated from camels in Hail region (Saudi Arabia) using massive sequencing of 16S rDNA. Bioinformation 16:602–610. doi:10.6026/97320630016602

Ankrah NYD, Chouaia B, Douglas E. 2018. The cost of metabolic interactions in symbioses between insects and bacteria with reduced genomes. MBio 9:1–15. doi: 10.1128/mBio.01433-18

Barradas PF, Lima C, Cardoso L, Amorim I, Gärtner F, Mesquita JR. 2020. Molecular evidence of *Hemolivia mauritanica*, *Ehrlichia spp.* and the endosymbiont *Candidatus* Midichloria mitochondrii in *Hyalomma aegyptium* infesting *Testudo graeca* tortoises from Doha, Qatar. animals. doi:10.3390/ani11010030

Brinkmann A, Hekimoǧlu O, Dinçer E, Hagedorn P, Nitsche A, Ergünay K. 2019. A cross-sectional screening by next-generation sequencing reveals *Rickettsia*, *Coxiella*, *Francisella*, *Borrelia*, *Babesia*, *Theileria* and *Hemolivia* species in ticks from Anatolia. Parasites and Vectors 12:1–13. doi:10.1186/s13071-018-3277-7

Buysse M, Duron O. 2021. Evidence that microbes identified as tick-borne pathogens are nutritional endosymbionts. Cell. doi: 10.1016/j.cell.2021.03.053

Cafiso A, Bazzocchi C, De Marco L, Opara MN, Sassera D, Plantard O. 2016. Molecular screening for *Midichloria* in hard and soft ticks reveals variable prevalence levels and bacterial loads in different tick species. Ticks Tick Borne Dis 7:1186–1192. doi:10.1016/j.ttbdis.2016.07.017

Cazalet C, Rusniok C, Brüggemann H, Zidane N, Magnier A, Ma L, Tichit M, Jarraud S, Bouchier C, Vandenesch F, Kunst F, Etienne J, Glaser P, Buchrieser C. 2004. Evidence in the *Legionella pneumophila* genome for exploitation of host cell functions and high genome plasticity. Nat Genet 36:1165–1173. doi:10.1038/ng1447

Collins NE, Liebenberg J, De Villiers EP, Brayton KA, Louw E, Pretorius A, Faber FE, Van Heerden H, Josemans A, Van Kleef M, Steyn HC, Van Strijp MF, Zweygarth E, Jongejan F, Maillard JC, Berthier D, Botha M, Joubert F, Corton CH, Thomson NR, Allsopp MT, Allsopp BA. 2005. The genome of the heartwater agent *Ehrlichia ruminantium* contains multiple tandem repeats of actively variable copy number. Proc Natl Acad Sci U S A 102:838–843. doi:10.1073/pnas.0406633102

Díaz-Sánchez S, Fernández AM, Habela MA, Calero-Bernal R, de Mera IGF, de la Fuente J. 2021. Microbial community of *Hyalomma lusitanicum* is dominated by *Francisella*-like endosymbiont. Ticks Tick Borne Dis 12. doi:10.1016/j.ttbdis.2020.101624

Dunning Hotopp JC, Lin M, Madupu R, Crabtree J, Angiuoli S V., Eisen J, Seshadri R, Ren Q, Wu M, Utterback TR, Smith S, Lewis M, Khouri H, Zhang C, Niu H, Lin Q, Ohashi N, Zhi N, Nelson W, Brinkac LM, Dodson RJ, Rosovitz MJ, Sundaram J, Daugherty SC, Davidsen T, Durkin AS, Gwinn M, Haft DH, Selengut JD, Sullivan SA, Zafar N, Zhou L, Benahmed F, Forberger H, Halpin R, Mulligan S, Robinson J, White O, Rikihisa Y, Tettelin H. 2006. Comparative genomics of emerging human ehrlichiosis agents. PLoS Genet 2:208–223. doi:10.1371/journal.pgen.0020021

Elbir H, Almathen F, Alhumam NA. 2019. A glimpse of the bacteriome of *Hyalomma dromedarii* ticks infesting camels reveals human Helicobacter pylori pathogen. J Infect Dev Ctries 13:1001–1012. doi:10.3855/JIDC.11604

Elbir H, Almathen F, Elnahas A. 2020. Low genetic diversity among *Francisella*-like endosymbionts within different genotypes of *Hyalomma dromedarii* ticks infesting camels in Saudi Arabia. Vet World 13:1462–1472. doi:10.14202/vetworld.2020.1462-1472

Epis S, Sassera D, Beninati T, Lo N, Beati L, Piesman J, Rinaldi L, McCoy KD, Torina A, Sacchi L, Clementi E, Genchi M, Magnino S, Bandi C. 2008. *Midichloria mitochondrii* is widespread in hard ticks (Ixodidae) and resides in the mitochondria of phylogenetically diverse species. Parasitology 135:485–494. doi:10.1017/S0031182007004052

Ghoneim NH, Abdel-Moein KA, Zaher HM. 2017. Molecular detection of *Francisella spp*. among ticks attached to camels in Egypt. Vector-Borne Zoonotic Dis 17:384–387. doi:10.1089/vbz.2016.2100

Ghosh S, Sela N, Kontsedalov S, Lebedev G, Haines LR, Ghanim M. 2020. An intranuclear *Sodalis*-like symbiont and *Spiroplasma* coinfect the carrot psyllid, *Bactericera trigonic*a (Hemiptera, Psylloidea). microorganisms 8. doi: 10.3390/microorganisms8050692

Gillespie JJ, Drisco TP, Verhoeve VI, Utsuki T, Husseneder C, Chouljenko VN, Azad AF, MacAluso KR. 2014. Genomic diversification in strains of *Rickettsia felis* isolated from different arthropods. Genome Biol Evol 7:35–56. doi:10.1093/gbe/evu262

Ivanov IN, Mitkova N, Reye AL, Hübschen JM, Vatcheva-Dobrevska RS, Dobreva EG, Kantardjiev T V., Muller CP. 2011. Detection of new *Francisella*-like tick endosymbionts in *Hyalomma spp.* and *Rhipicephalus spp.* (Acari: Ixodidae) from Bulgaria. Appl Environ Microbiol 77:5562–5565. doi:10.1128/AEM.02934-10

Kurtti TJ, Felsheim RF, Burkhardt NY, Oliver JD, Heu CC, Munderloh UG. 2015. *Rickettsia buchneri* sp. nov., a rickettsial endosymbiont of the blacklegged tick *Ixodes scapularis*. Int J Syst Evol Microbiol 65:965–970. doi:10.1099/ijs.0.000047

Larsson P, Oyston PCF, Chain P, Chu MC, Duffield M, Fuxelius HH, Garcia E, Hälltorp G, Johansson D, Isherwood KE, Karp PD, Larsson E, Liu Y, Michell S, Prior J, Prior R, Malfatti S, Sjöstedt A, Svensson K, Thompson N, Vergez L, Wagg JK, Wren BW, Lindler LE, Andersson SGE, Forsman M, Titball RW. 2005. The complete genome sequence of *Francisella tularensis*, the causative agent of tularemia. Nat Genet 37:153–159. doi:10.1038/ng1499

Lin M, Zhang C, Gibson K, Rikihisa Y. 2009. Analysis of complete genome sequence of *Neorickettsia risticii*: Causative agent of Potomac horse fever. Nucleic Acids Res 37:6076–6091. doi:10.1093/nar/gkp642

Lopes de Carvalho I, Toledo A, Carvalho CL, Barandika JF, Respicio-Kingry LB, Garcia-Amil C, García-Pérez AL, Olmeda AS, Zé-Zé L, Petersen JM, Anda P, Núncio MS, Escudero R. 2016. *Francisella* species in ticks and animals, Iberian Peninsula. Ticks Tick Borne Dis 7:159–165. doi:10.1016/j.ttbdis.2015.10.009

Luo J, Liu MX, Ren QY, Chen Z, Tian ZC, Hao JW, Wu F, Liu XC, Luo JX, Yin H, Wang H, Liu GY. 2017. Micropathogen community analysis in *Hyalomma rufipes* via high-throughput sequencing of small RNAs. Front Cell Infect Microbiol 7:1–12. doi:10.3389/fcimb.2017.00374

Perveen N, Muzaffar S Bin, Vijayan R, Al-Deeb MA. 2020. Microbial communities associated with the camel tick, Hyalomma dromedarii: 16S rRNA gene-based analysis. Sci Rep 10:1–11. doi:10.1038/s41598-020-74116-7

Ravi A, Ereqat S, Al-Jawabreh A, Abdeen Z, Shamma OA, Hall H, Pallen MJ, Nasereddin A. 2018. Metagenomic profiling of ticks: Identification of novel rickettsial genomes and detection of tick borne canine parvovirus. bioRxiv 1–19. doi:10.1101/407510

Rio RVM, Symula RE, Wang J, Lohs C, Wu Y neng, Snyder AK, Bjornson RD, Oshima K, Biehl BS, Perna NT, Hattori M, Aksoy S. 2012. Insight into the transmission biology and species-specific functional capabilities of Tsetse (Diptera: Glossinidae) obligate symbiont *Wigglesworthia*. MBio 3:1–13. doi:10.1128/mBio.00240-11

Sait M, Aitchison K, Wheelhouse N, Wilson K, Lainson FA, Longbottom D, Smith DGE. 2013. Genome sequence of *Lawsonia intracellularis* strain N343, isolated from a sow with hemorrhagic proliferative enteropathy. Genome Announc 1:1–2. doi:10.1128/genomeA.00027-13

Sassera D, Lo N, Epis S, D’Auria G, Montagna M, Comandatore F, Horner D, Peretó J, Luciano AM, Franciosi F, Ferri E, Crotti E, Bazzocchi C, Daffonchio D, Sacchi L, Moya A, Latorre A, Bandi C. 2011. Phylogenomic evidence for the presence of a flagellum and cbb3 oxidase in the free-living mitochondrial ancestor. Mol Biol Evol 28:3285–3296. doi:10.1093/molbev/msr159

Seshadri R, Paulsen IT, Eisen JA, Read TD, Nelson KE, Nelson WC, Ward NL, Tettelin H, Davidsen TM, Beanan MJ, Deboy RT, Daugherty SC, Brinkac LM, Madupu R, Dodson RJ, Khouri HM, Lee KH, Carty HA, Scanlan D, Heinzen RA, Thompson HA, Samuel JE, Fraser CM, Heidelberg JF. 2003. Complete genome sequence of the Q-fever pathogen *Coxiella burnetii*. Proc Natl Acad Sci U S A 100:5455–5460. doi:10.1073/pnas.0931379100

Sjödin A, Svensson K, Öhrman C, Ahlinder J, Lindgren P, Duodu S, Johansson A, Colquhoun DJ, Larsson P, Forsman M. 2012. Genome characterisation of the genus *Francisella* reveals insight into similar evolutionary paths in pathogens of mammals and fish. BMC Genomics 13. doi:10.1186/1471-2164-13-268

Szigeti A, Kreizinger Z, Hornok S, Abichu G, Gyuranecz M. 2014. Detection of *Francisella*-like endosymbiont in *Hyalomma rufipes* from Ethiopia. Ticks Tick Borne Dis 5:818–820. doi:10.1016/j.ttbdis.2014.06.002

Visser ES, McGuire TC, Palmer GH, Davis WC, Shkap V, Pipano E, Knowles DP. 1992. The *Anaplasma marginale msp5* gene encodes a 19-kilodalton protein conserved in all recognized *Anaplasma* species. Infect Immun 60:5139–5144. doi:10.1128/iai.60.12.5139-5144.1992

Wang Y, Mao L, Sun Y, Wang Z, Zhang Jiayong, Zhang Jibo, Peng Y, Xia L. 2018. A novel *Francisella*-like endosymbiont in *Haemaphysalis longicornis* and *Hyalomma asiaticum*, China. Vector-Borne Zoonotic Dis 18:669–676. doi:10.1089/vbz.2017.2252

